# ADAPTATION AGAINST LONG-TERM PROTEIN DEPRIVATION TRADES OFF WITH IMMUNOCOMPETENCE AND THE ABILITY TO SURVIVE PATHOGENIC INFECTIONS

**DOI:** 10.1101/2025.09.10.675413

**Authors:** Saubhik Sarkar, Dipendra Nath Basu, Simran Sethi, Bodhisatta Nandy, Imroze Khan

**Affiliations:** Department of Biology, Ashoka University, Plot No. 2, Rajiv Gandhi Education City, National Capital Region, P.O. Rai, Sonepat, Haryana-131029, India; Indian Institute of Science Education and Research Berhampur, Ganjam, Berhampur-760010, Odisha, India

**Keywords:** Experimental evolution, Protein limitations, Sexual dimorphism, Trade-offs, Transcriptomics

## Abstract

Insufficient protein intake, leading to malnutrition, is a major global health concern that compromises the immune system and increases susceptibility to diseases. In scenarios where protein availability is constrained, organisms may experience strong selection to efficiently allocate their resources between immunity vs other energy-demanding processes, such as reproduction, resulting in evolutionary trade-offs. Additionally, in many species, protein deficiency has a more significant impact on female reproduction than on males, potentially leading to pronounced sexually dimorphic trade-offs involving immunity and infection outcomes. However, there are no experiments to test these possibilities. In this work, we demonstrate that in *Drosophila melanogaster* populations selected for increased early-life reproduction under protein limitations, evolved virgin females indeed suffered a greater reduction in their resistance to the pathogen *Providencia rettgeri* and showed lower survival following infection than males, corroborating our expectations. However, mating resulted in a loss of sexually dimorphic infection outcomes, causing both sexes to exhibit nearly identical infection costs and reduced infection tolerance compared to their ancestral, unselected populations. Moreover, several immune components, including the Toll- and IMD-mediated inducible immune pathways, were either less upregulated or more downregulated in the selected flies, which may contribute to their heightened susceptibility to pathogens. Downregulation in key metabolic pathways and genes related to phagocytosis, melanisation and ROS-mediated defence after infection in selected flies can also be associated with their increased pathogen vulnerability. Taken together, our work thus reveals the reproductive status- and sex-specific plasticity of immune investments and post–infection health in response to evolutionary constraints under chronic protein deprivation.

## INTRODUCTION

During the early stages of life, dietary protein is essential for both growth and maintenance, whereas after reaching sexual maturity, it plays a pivotal role in supporting reproduction (Kaushik et al., 1995; Ma et al., 2022). Previous research indicates that variations in dietary protein impact several reproductive aspects, including gonadal development, gametogenesis, and the quantity and quality of eggs, as well as fertilisation rates and offspring viability (Herring et al., 2018; Radhakrishnan et al., 2020). Additionally, organisms across various taxa experience reduced longevity and produce lower-quality offspring when maintained on a low-protein diet (Simpson and Raubenheimer, 2009; Lee 2015). For instance, juvenile protein restriction has been linked to prolonged developmental time in *Drosophila melanogaster* (Krittika et al., 2019), decreased adult lifespan in *Telostylinus angusticollis* (Runagall-McNaull et al., 2015), and slower increase in body weight in growing mice *Mus musculus*, resulting in low body mass index (Madhura et al., 2023). Furthermore, adult protein deprivation causes underdeveloped ovaries and infertility in *T. angusticollis* females (Adler et al., 2013), weight loss, and notable brain structural changes alongside behavioural defects in adult mice (Wu et al., 2021; Lukoyanov and Andrade, 2000). Reduced performance in various fitness measures due to dietary protein limitations may thus exert strong selection pressure, prompting organisms to evolve compensatory mechanisms to mitigate fitness costs.

Indeed, a recent study reveals that when laboratory-adapted populations of fruit flies, *D. melanogaster* experience chronic protein deprivation, by limiting live-yeast supplementation before reproduction (Dasgupta et al., 2022), females can rapidly evolve to enhance early-life fecundity by increasing protein content at eclosion and reallocating lipids to reproduction (Dasgupta et al., 2024). However, this early-life resource allocation towards maximising reproductive success may trade off with life-history traits that also rely heavily on protein availability (Cotter et al., 2011) and lipid reserve (Wrońska et al., 2023). For instance, since immune responses are costly (Moret and Schmid-Hempel, 2000), they provide a classic example of such trade-offs where organisms prioritising reproduction may lack sufficient protein reserves to mount effective immune responses, making them more susceptible to pathogens (Demas et al., 2012; Schwenke et al., 2016). In fact, this possibility has been corroborated by several studies, where lower protein availability has reduced multiple components of cellular and humoral immunity, as well as survival after bacterial infection, in *Spodoptera littoralis* caterpillars, fruit flies, and mice (Meshrif et al., 2022; Taylor et al., 2013; Povey et al., 2014). Conversely, infected animals prefer food with a higher protein-to-carbohydrate ratio, which enhances hemocyte circulation, antibacterial activity, and resistance to bacterial infections (Povey et al., 2009). This highlights the crucial role that dietary protein may have in shaping immune responses. Additionally, previous studies on fruit flies suggest that activating the immune response can affect anabolic lipid metabolism by reducing lipid storage and increasing phospholipid production, which are crucial for synthesising and releasing antimicrobial peptides (Martinez et al., 2020). If there are significant demands for lipid storage to enhance reproduction, such a situation may lead to trade-offs with immunity and infection responses (Moghadam et al., 2015).

Life history and sexual selection theories suggest that the strength of such trade-offs may also differ between sexes, driven by inherent sex differences in immune investments, thus leading to sexually dimorphic infection outcomes (Zuk and McKean, 1996; Khan and Prasad, 2011; Belmonte et al., 2020). Typically, females are expected to invest in a long, healthy lifespan to maximise lifetime reproductive output, and hence, may invest more in immunity, thereby being less susceptible to infections (Rolff 2002; Zuk 2009). Primarily due to sexual selection, males, on the other hand, may prioritise reproductive effort and invest less in immunity, and hence be more susceptible to infection (Bonduriansky et al., 2008; Adler and Bonduriansky, 2014). However, a low-protein environment may bring further complications. For example, if selection primarily works on fecundity, under chronic protein deprivation, females may be more strongly selected to reduce investment in immunity for reproductive gains (Fedorka et al. 2005; Rodrigues et al., 2021). Indeed, a prior study involving fruit flies indicated that restricted access to dietary yeast significantly compromised immune function in females but not in males (McKean and Nunney, 2005). This effect may become more pronounced following mating, particularly in females actively involved in egg production (Short and Lazzaro, 2010). In contrast, males might be less affected, as their reproductive success is not as dependent on protein intake as females’ is, but rather on secondary sexual traits (Lee et al., 2013), making them less responsive to dietary changes.

In this work, we tested these possibilities using replicate populations of the same outbred *D. melanogaster* populations as described in Dasgupta et al. (2022). These populations have been selected for survival and reproduction under chronic live-yeast deprivation for over 100 generations and have evolved increased early-life fecundity as an adaptive phenotype. We began by infecting them with the Gram-negative bacterial pathogen *Providencia rettgeri* (Pr) across sexes and mating status. By and large, we found that the selected flies succumbed to infection more frequently than those from the control population; however, the ability to counter pathogen burden and the overall infection outcome varied widely across infection dose, sex, and mating status of the flies. For example, virgin female flies from the selected regime showed lower resistance to infection than control flies, resulting in consistently higher post–infection mortality, regardless of the infection dose. By contrast, the effects on virgin male flies from selected populations were lower than those on females. Upon mating, selected flies from both sexes became equally vulnerable to infection, suggesting a loss of observed sexual dimorphism. We also performed transcriptomic analyses that revealed lower upregulation of inducible immune components (e.g., receptors, regulators, and effectors of IMD and Toll immune pathways) and downregulation of phagocytosis, melanisation and ROS-mediated defense in selected flies, corroborating their increased susceptibility to pathogens. Taken together, our findings provide rare insights into how the evolutionary background of chronic protein deprivation may influence the design features of the immune system, with strong implications for responses to newly emerging infections.

## MATERIALS AND METHODS

### Fly populations and the experimental evolution paradigm

For this study, we used replicated *Drosophila melanogaster* populations adapted to a chronic dietary protein deprivation regime (See Dasgupta et al., 2022 for details). Briefly, the selected regime (YLB) was derived from the control populations (BL) and maintained in a 14-day discrete generation cycle. On the 12^th^ day after egg collection, we transferred the newly emerged adult flies from the control regime to plexiglass cages (dimension of 23cm × 20cm × 15 cm) with a food plate smeared with *ad-libitum* live yeast as a protein supplementation just before reproduction, whereas deprived the selected flies of live yeast to impose severe protein deficiency for over ~120 generations. Although both regimes had four replicate populations, we only used three randomly chosen populations from each regime for logistical reasons. To generate standardised experimental flies, we reared replicate blocks of control and selected flies with live yeast supplementation for one generation, thereby minimising parental and environmental effects (Sarkar et al., 2025) and collected experimental flies in the subsequent generation. For each replicate block, we collected both adult female and male flies and held them as virgins at a density of 15 flies/vial. We also allowed a set of flies to mate, and after 18 hours of mating, separated females and males in separate vials. For the assays, we used age-matched 3–4-days-old (post–eclosion) female and male flies. We infected virgin and mated flies on separate days due to logistical reasons.

### Bacterial infection and assays

We used a natural, opportunistic fly pathogen, the Gram-negative bacterium *Providencia rettgeri*, to induce septic infection in our experimental flies, following a protocol described in Sarkar and coworkers (2025). Briefly, for each sex and mating status, we pricked individual flies into their thoracic region using a 0.1 mm minuten pin (Fine science tool) dipped in bacterial suspension adjusted to 3 different concentrations, namely 1, 20, and 40 OD (corresponding to approximately 27.75, 83.25 and 134.79 cells respectively, measured at 600 nm, originally derived from a 10 ml of overnight-grown 1 OD culture of *P. rettgeri*) (n=75 flies/ infection dose/ sex/ mating status/ selection regime/ block). We also pricked flies with sterile phosphate-buffered saline (1X) as a procedural control (or sham infection). After infection (or sham-infection), we redistributed them in food vials in a batch of 15 individual flies, resulting in five independent replicate vials for each experimental treatment.

Following this, we randomly removed three flies from each vial and individually measured their bacterial loads 18 hours post–infection—i.e., the time point at which mortality began (i.e., the onset of virulence manifestation; Seal et al., 2025). As described in Sarkar and coworkers (2025), we extracted the whole-body homogenate of each fly and plated them on Luria-Agar plates to count the colony-forming units (CFUs) (Siva-Jothy et al., 2018). To analyse the bacterial load data, we first pseudo-log-transformed the CFUs and then used a generalised linear mixed-effects model, with infection dose, selection regime and sex as fixed effects and vial identity nested within the replicate blocks as a random effect (Model: Log bacterial load~ Infection dose × Selection regime × Mating status × Sex + 1|Block/Vial, family=negative binomial, using “glm.nb” function in glmmTMB package). However, due to multiple two and three-way interactions, we next analysed sexes and mating status separately to clearly distinguish the effect of selection (Model: Log bacterial load~ Infection dose [ID] × Selection regime [SR] + 1|Block/Vial, family=negative binomial; using glm.nb function in glmmTMB package) (Bates et al. 2014), followed by a post hoc test with Tukey’s adjustment to compare between regimes across different infection doses.

For the remaining 12 flies within each vial, we scored the number of dead flies every three hours for the first three days, followed by every six hours for the next two days. To analyse the survival data, we fitted a mixed-effects Cox regression model, using infection dose, selection regime, mating status, and sex as fixed effects, and vial identity nested within replicate blocks as a random effect (Post–infection survival ~ Infection dose × Selection regime × Mating status × Sex + 1| Block/ Vial, using “coxme” function in survminer package) (Therneau T, 2024). Similar to bacterial load analysis, we found multiple two- and three-way interactions here as well. We thus analysed sexes and mating status separately to clearly distinguish the effect of selection on post-infection survival (Post–infection survival ~ Infection dose × Selection regime + 1| Block/ Vial). Additionally, we quantified the level of susceptibility of infected flies across infection treatments by estimating hazard ratio (HR) of deaths occurring in each dose of *P. rettgeri* infected flies vs. sham-infected flies across sex, mating status and selection regimes (Khan et al., 2017). Note that since we did not find any mortality in the sham-infected individuals, we have included one dummy mortality in each vial containing sham-infected flies to calculate the hazard ratio of Pr-infected vs sham-infected treatments. Hazard ratios significantly greater than one indicated a higher mortality risk in the Pr-infected groups than in sham-infected ones.

We also quantified the level of susceptibility in Pr-infected flies by estimating hazard ratios of deaths occurring in selected vs. control flies across infection doses, mating status and sexes. We analysed the data using a generalised linear mixed-effects model, with infection dose, mating status, and sex as fixed effects, and vial identity nested within replicate blocks as a random effect (Model: Hazard ratio ~ Infection dose × Mating status × Sex [S] + 1|Block/Vial, family=negative binomial), followed by a post hoc test with Tukey’s adjustment).

Note that our experimental design enabled us to collect paired datasets for post–infection host health, estimated as the hazard ratio, and bacterial load for each replicate vial across infection doses and selection regimes. We thus next analysed changes in hazard ratio as a response to variations in bacterial load using a generalised linear model, with bacterial load as a continuous covariate and selection regime as fixed effects for each sex and mating status separately (Model: Hazard ratio ~ Log average bacterial load × selection regime, family = negative binomial). A significant interaction between bacterial load and selection regime would indicate divergence in the rate at which health declines as a function of increasing bacterial load across different selection regimes, serving as a measure of infection tolerance (see Gupta and Vale, 2017; Seal et al., 2021).

### Transcriptomic analyses

To identify the genetic basis for infection response to selection under chronic protein deprivation, we performed whole-body transcriptome analyses of mated female flies from control and selected regime after *Providencia rettgeri* infection (and sham-infection as a procedural control) (n= 10 females pooled together from each infection treatment/selection regime/ replicate/ block). Attributed to the virulence manifestation of *P. rettgeri*, we sampled flies at 18 hours post–infection. Pooled individuals were snap-frozen in liquid nitrogen and stored at −80°C until further steps. We used the Aurum total RNA mini kit from Biorad to extract RNA following the manufacturer’s protocol and outsourced the samples for sequencing. The quantity and quality of extracted RNA was checked on a Qubit 4.0 fluorometer (Thermofisher #Q33238) using an HS RNA assay kit (Thermofisher #Q32851) and on TapeStation using HS RNA ScreenTape (Agilent), respectively. The libraries were prepared using TruSeq® Stranded Total RNA kit (Illumina #15032618, Illumina #20020596) post–poly-A enrichment. The sequencing was performed on an Illumina Novaseq 6000 platform using a 150 bp paired-end chemistry.

Next, we passed the raw reads through FastQC and MultiQC to check the quality parameter summaries. We then used Cutadapt 4.6 to remove adapter contamination using standard Illumina adapter sequences (forward adapter sequence: AGATCGGAAGAGCACACGTCTGAACTCCAGTCA, reverse adapter sequence: AGATCGGAAGAGCGTCGTGTAGGGAAAGAGTGT along with poly-A tail removal). During adapter trimming, we discarded reads with N base counts of more than 10% of their length and more than 10 low-quality bases and restricted the final read lengths to a minimum of 70 base pairs for the subsequent analyses. We then created a genomic index using the *D. melanogaster* reference genome GCF_000001215.4 and mapped all the reads using Hisat2, allowing for up to three mismatches per read, along with strand information (Hoskins et al., 2015). We used stringtie 2 for sorting bam output files. Subsequently we used HtSeq2 to obtain the raw read count. After normalizing raw reads, we estimated the differential gene expressions (DEGs) using the R package “DESeq2” and used “pheatmap” and “RColorBrewer” packages to visualize the expression profiles of all the DEGs by a pooled-population heatmap based on the z-score of normalized read counts. To this end, we counted the common vs. unique sets of up-and down-regulated DEGs as a response to infection treatment, with contrasting expressions between regimes and visualized them using the R-package “ComplexUpset”. We performed a principal component analysis based on the normalized count data of all the DEGs from the selected and control regimes to examine the effect of the interaction between regime and infection on the expression changes. We further used these GO terms to perform pathway enrichment for up and down-regulated DEGs separately, using String protein database references in Cytoscape 3.0 and visualised only the KEGG pathways using the R-package “ggplot2”. Subsequently, to understand the possible role of immune responses, we categorised DEGs with known immunological roles into six broad functional categories, (a) immune receptors, (b) immune regulators and (c) immune effectors driving inducible immunity and markers for (d) melanization and coagulation, (e) encapsulation and phagocytosis, and (f) ROS mediated defence (Westlake et al., 2024), and performed MANOVA to find the effect infection treatment and selection regime. To further explore the associations, we performed a canonical variant analysis (CVA) to obtain a metagene expression profile, separating the effects of infection and selection regimes in each immune and reproductive gene category. We corroborated the graphical representation of gene expression profiles across infection and selection regimes with statistical differences found in infection, selection regime and their interactions from MANOVA tests. We used R version 4.2.3 for all analyses (Therneau, 2014).

## RESULTS

### Adaptation against protein restriction trade-offs with post–infection survival

A mixed-effects Cox regression model on post–infection survival data revealed the main effects of selection regime, sex, infection dose, mating status, and multiple two- and three-way interactions (**Tables S1**). To disentangle the effect of the selection regime more clearly, we thus analysed the mating status and sexes separately (**Tables S2–S4**). Overall, the selection regime had a significant impact on both sexes, with evolved populations showing higher susceptibility to pathogenic infections (**Figure 1A–1B**; **Tables S2–S4**). We also found significant interactions between selection regimes and infection doses in virgin flies but not in mated flies (**Figure 1A, 1B**; **Table S2**). In virgin females, the impact of the selection regime was consistent across all infection doses and became stronger with higher doses of infection (e.g., comparing the effect size in **Table S3**). In contrast, the effect in males was detected only at an infection dose of 40 OD (**Figure 1A**; **Table S3**).

**Figure 1:**
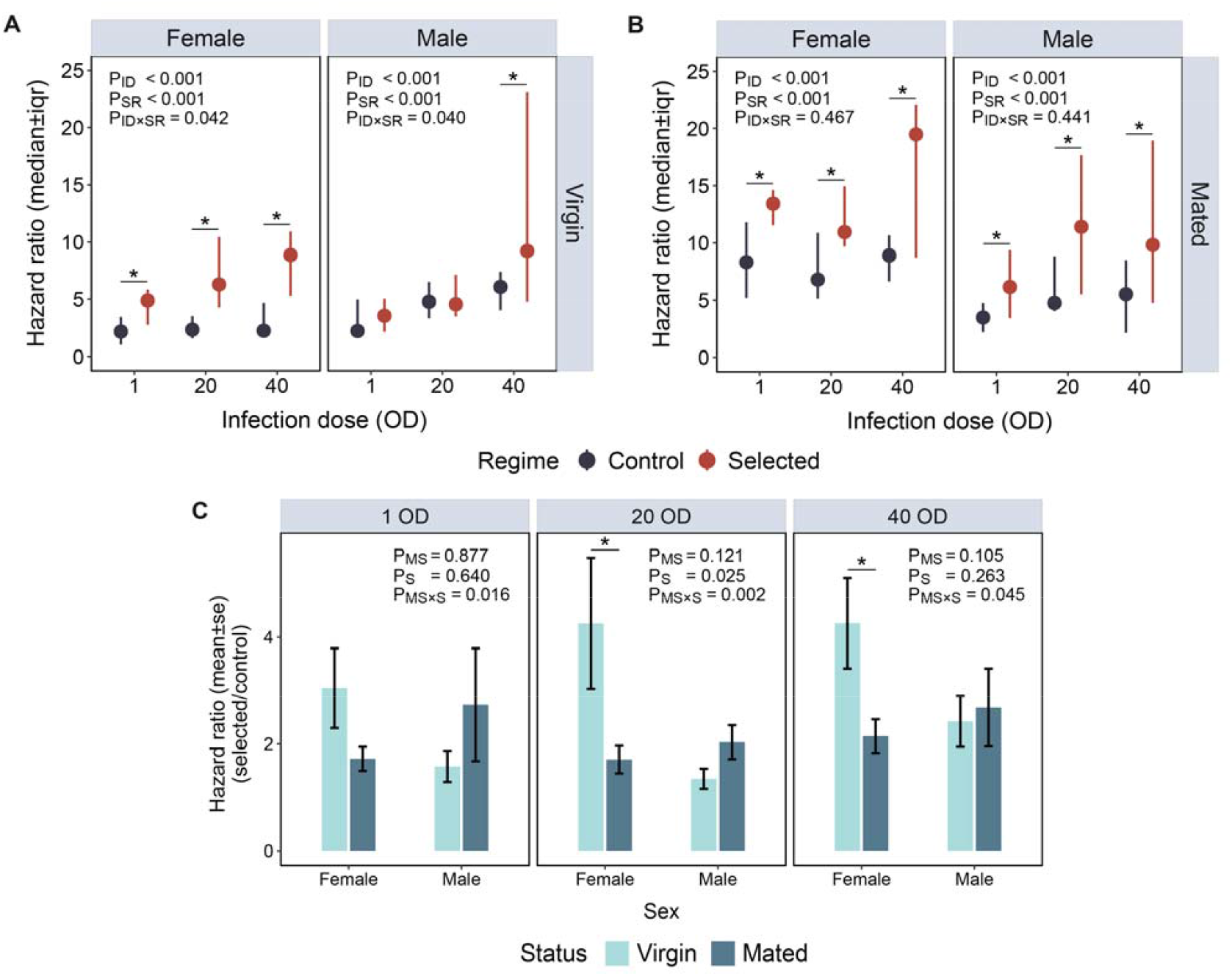
Post–infection survival response of control and selected regime, adapted to chronic protein malnourishment. Hazard ratio for the survival response 120 h post–infection of **(A)** virgin and **(B)** mated flies of both sexes against *Providencia rettgeri* infection in control vs selected regimes. Each dot represents the median±iqr hazard ratio of five replicate vials of flies that were assayed for their survival within each experimental treatment (i.e., n= 12 flies/ replicate vials/ infection dose/ sex/ selection regime/ block). The p values are obtained from a mixed-effect cox regression (coxme) model (Model: *Post*–*infection survival ~ Infection dose [ID] × Selection regime [SR] +1*| *Replicate population/ Vial*). The pairwise comparisons between regimes at individual doses are obtained from a post hoc test with Tukey’s adjustment, where only statistically significant differences between regimes are indicated with an asterisk (*) (i.e., p < 0.05). In panel A-B, infection dose is designated by ID, selection regime is designated by SR, and the interaction term is indicated by ID×SR. **(C)** Hazard ratio measured for selected regime as compared to controls against *Providencia rettgeri* infection in virgin vs mated status of both sexes. The p values are obtained from a generalized linear mixed effect (GLMM) model (Model: *Hazard ratio ~ Mating status [MS] × Sex [S] + 1*|*Block/Vial, family=negative binomial*). The pairwise comparisons between mating status for each sex are obtained from a post hoc test with Tukey’s adjustment, where only statistically significant differences between mating status are indicated with an asterisk (*) (i.e., p < 0.05). In the graph, mating status is designated by MS, sex is designated by S, and the interaction term is indicated by MS×S.

Given the high variance across replicate populations (or blocks) within each selection regimes (**Table S2**), we also analysed each block separately. We found that the effect of the selection regime was more consistent in virgin females, such that in two out of three replicate blocks (Block 1 and 3; Block 2 showed a weak trend), selection regime had significant impacts (**Figure S1A**; **Table S5**), whereas only one replicate block (Block 1) drove the effects in virgin males (**Figure 1A, S1A**; **Table S3, S6**). Upon mating, the impact of the selection regime became more consistent across both sexes (**Figure 1B**; **Table S4**). More specifically, two out of three blocks exhibited a significant decline in post-infection survival in both sexes (Females: Block 2, 3; Males: Block 2, 3) (**Figure S1B**; **Table S7–S8**). Here, it is important to note that, when considering both virgin and mated conditions, females exhibited selection effects in all the three blocks (i.e., Blocks 1–3). In comparison, males exhibited effects in just two blocks, despite pooling the observations across mating status.

Moreover, when we compared the effect size of selection treatment, estimated as the hazard ratio of post– infection survival between selected and control flies, we found that the relative decline in post–infection health was higher in virgin females than in virgin males, suggesting sex-specific variations in trade-offs (**Figure 1C, Table S9–S10**). However, such effects were not observed in mated flies. Moreover, the effect of selection was weaker in mated females than in their virgin counterparts, which was possibly driven by basally higher infection costs associated with mating in control females (**Figure 1C, Table S10**). On the contrary, we did not find a significant effect of mating on selection regime in males, as both virgin and mated males showed similar effects of selection regimes on post–infection survival (**Figure 1C, Table S10**).

### Pathogen load did not explain the variation in post–infection survival consistently

A generalised linear mixed-effects model on bacterial load revealed the main effects of selection regime, sex, infection dose, mating status, and multiple two- and three-way interactions (**Tables S11**). To understand the effect of the selection regime more clearly, we thus analysed the mating status and sexes separately (**Tables S12–S14**). Virgin females from selected regimes were less efficient in clearing the bacterial cells than their control counterparts (**Figure 2A, Table S12–S13**), which explained their higher vulnerability to infection (**Figure 1A**). However, we did not find any variation in their tolerance, as females from both regimes suffered a similar loss of post–infection health, estimated as the hazard ratio of infected vs. sham-infected individuals, with increasing bacterial load (**Figure 2C, Table S15**). In contrast, although virgin males from both selected and control regimes showed no difference in their bacterial load (**Figure 2A, Table S12–S13**), selected males showed lower tolerance to infection than controls (i.e., faster rate of post–infection health loss with increasing bacterial load) (**Figure 2C, Table S15**). However, this can be attributed to the disproportionately higher hazard ratios observed in selected males in Block 1 (**Figure S1A, Table S6**). We did not find any difference in bacterial load (**Figure 2B, Table S12, S14**) and pathogen tolerance (**Figure 2D, Table S16**) between control and selected individuals after mating, regardless of their sex. Interestingly, when compared across sexes, mated females from the selected regime appeared to tolerate the infection better than their male counterparts, as they experienced a slower decline in post–infection health with increasing bacterial load (GLM: Hazard ratio ~ Log average bacterial load [BL] × Sex [S], family = negative binomial; P_BL_ < 0.001, P_S_ < 0.001, P_BL × S_ < 0.001). However, the post–infection mortality of selected females was already higher than that of males, even at a lower *P. rettgeri* dose, suggesting already poor health, which could not worsen further (**Figure 2D**).

**Figure 2:**
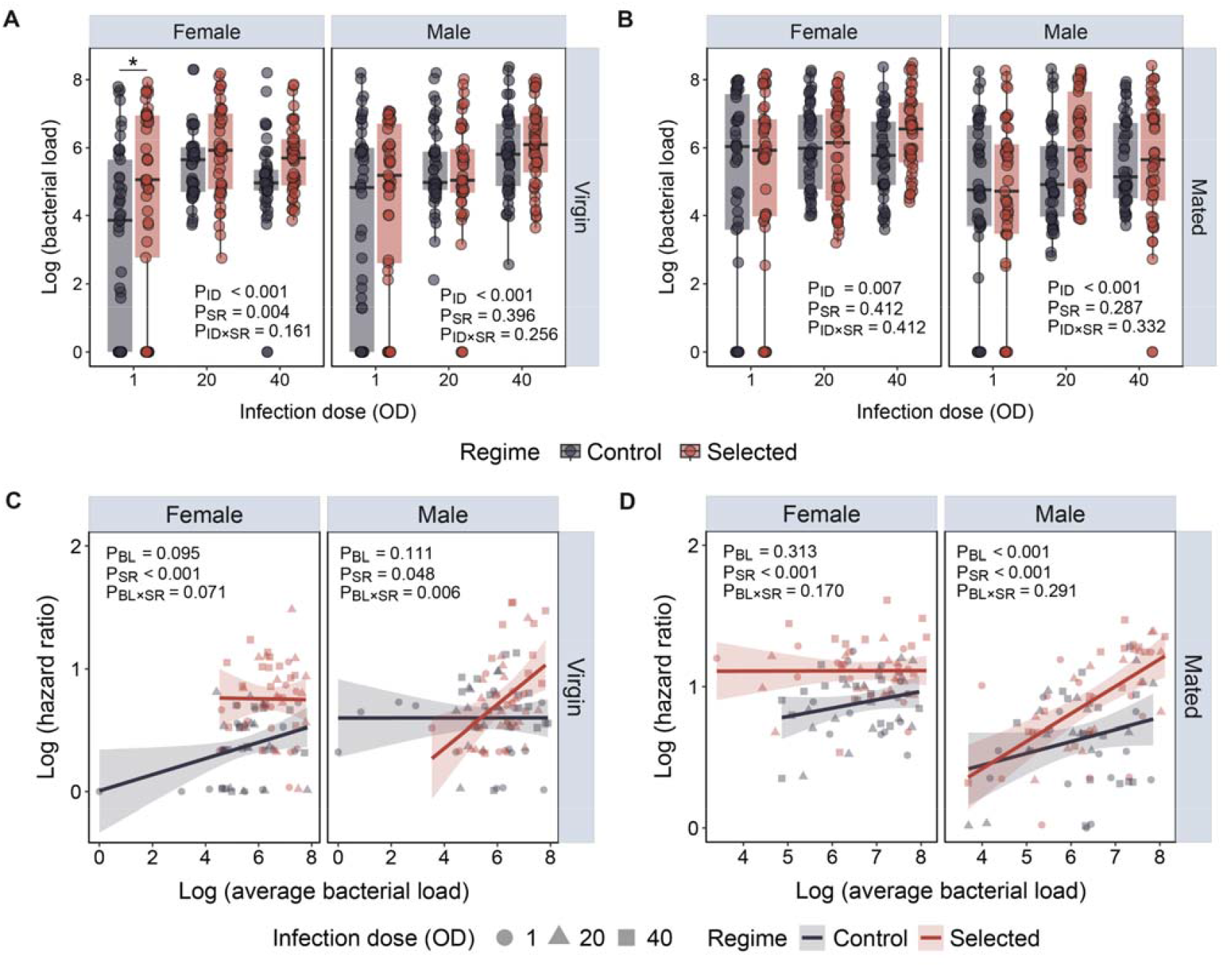
Bacterial load and infection tolerance of control and selected regime, adapted to chronic protein malnourishment. Bacterial load measured for **(A)** virgin and **(B)** mated flies of both sexes against *Providencia rettgeri* infection in control vs selected regimes at 18 h post–infection. Each dot represents a log-transformed CFU (colony forming unit) from individual flies sampled from each of the 5 replicated vials across treatments (n= 3 fliles/ replicate vials/ infection dose/ sex/ selection regime/ block). The p-values are obtained from a generalised linear mixed effect model (GLMM) (Model: *Log bacterial load ~ Infection dose [ID] × Selection regime [SR]* + 1|Replicate population/Vial, *family=negative binomial*). The pairwise comparisons between regimes at individual doses are obtained from a post hoc test with Tukey’s adjustment, where only statistically significant differences between regimes at each infection dose are indicated with an asterisk (*) (i.e., p < 0.05). In panel A-B, infection dose is designated by ID, selection regime is designated by SR, and the interaction term is indicated by ID×SR. Reaction norm of post–infection health estimated as hazard ratios vs the corresponding bacterial load (i.e., variation in post–infection health with changes in pathogen burden; or infection tolerance) in **(C)** virgin and **(D)** mated flies of both sexes against *Providencia rettgeri* infection, pooled across selection regimes for each block. Each shape (circle=1 OD, triangle=20 OD, square =40 OD) represents the log-transformed hazard ratio and the correlated estimate of pseudo-log-transformed bacterial load derived from each replicate vial. The p values are obtained from a generalized linear model (GLM) (Model: *Log hazard ratio ~ Log average bacterial load [BL] × Selection regime [SR], family=negative binomial*). A significant interaction between bacterial load and selection regime indicates variation in infection tolerance across regimes. In panel C-D, bacterial load is designated by BL, selection regime is designated by SR, and the interaction term is indicated by BL×SR.

### Adaptation to chronic live yeast deprivation leads to divergent gene expression patterns across immune components in response to infection

To gain molecular insights into the differential post-infection survival driven by chronic live yeast deprivation, we conducted transcriptome analyses and found that major variance in gene expression pattern was a function of the interaction between infection status and selection regime of the flies (**Figure 3A**). Subsequently, to disentangle the changes in immune response affected by adaptation to protein deprivation, we identified differentially expressed genes (DEGs) by comparing the gene expression of sham-infected and *Pr* infected flies from control and selected regimes separately. We obtained a higher number of differentially expressed genes (DEGs) in the selected flies (N = 892) than in their control counterparts (N = 390) (**Figure 3B**). Among these DEGs, 293 were common to both control and selected flies, comprising gene molecules that consistently showed an expression change in the presence of infection, regardless of their selection regime.

**Figure 3:**
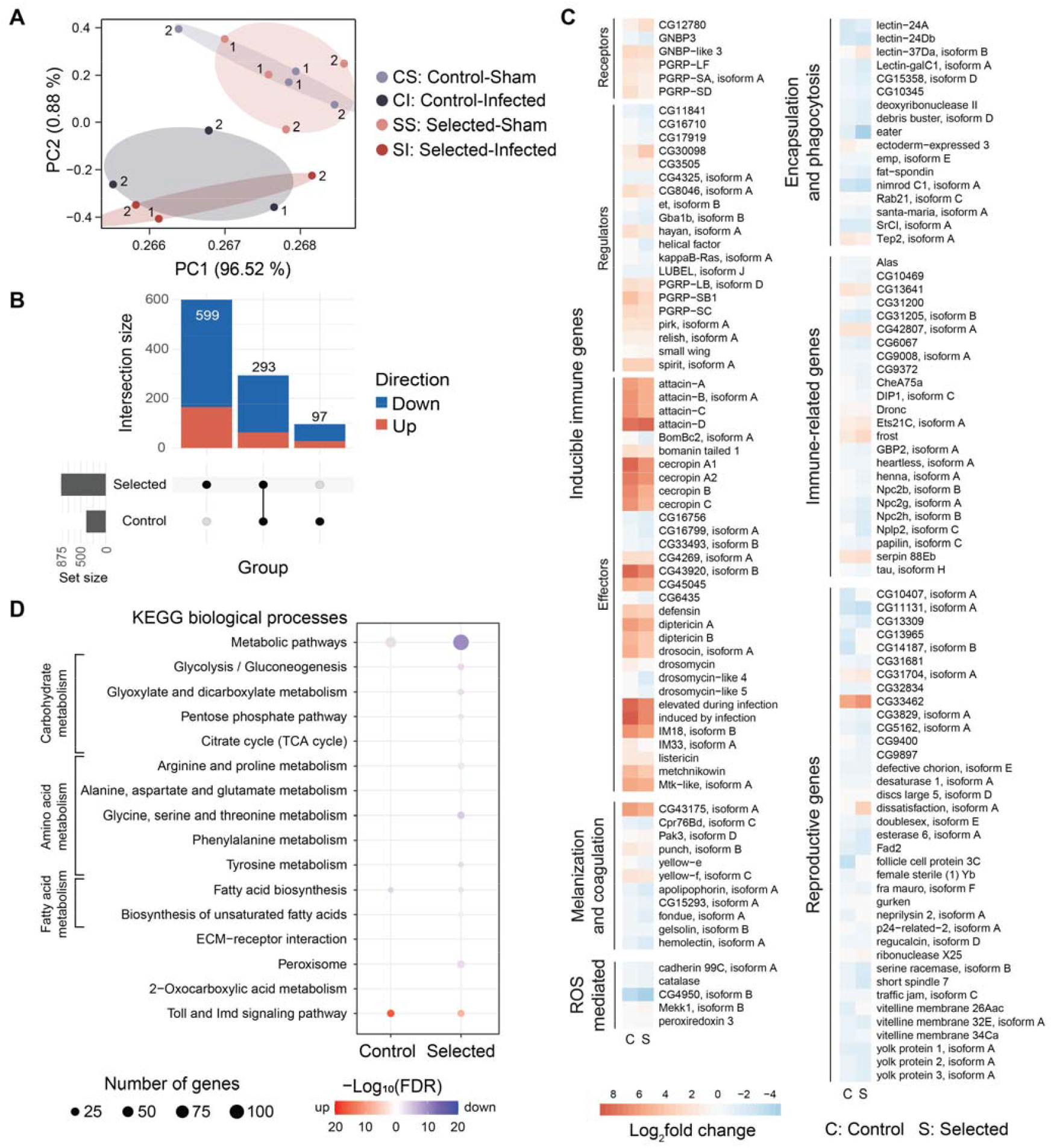
Transcriptomics analyses in control and selected flies adapted to chronic protein malnourishment after infection. **(A)** Principal component analysis on differentially expressed genes in control vs selected regimes after infection treatments (e.g., sham infection vs *Providencia rettgeri* infection). Numbers denote replicate blocks. Each data point is further colour coded respective to their selection regime and experimental treatments **(B)** Upset plot showing the total number of common and unique differentially expressed genes (DEGs) between control vs selected regimes. Red denotes proportion of upregulated genes and blue denotes proportion of downregulated genes due to infection. **(C)** Heatmaps of differentially expressed genes, involved in different immunological functions (divided into broad functional categories: a. immune receptors, c. immune regulators, and c. immune effectors driving inducible immune response, d. melanization and coagulation, e. ROS mediated response, f. encapsulation and phagocytosis) and reproduction, after infection treatments (e.g., sham infection vs *Providencia rettgeri* infection) in control vs selected regimes. The heatmap is based on Z-score estimated from normalized read counts; **(D)** Enrichment of major metabolic pathways that are either differentially upregulated (red) or downregulated (blue) post-infection in control vs selected regimes.

Conversely, there were 97 and 599 DEGs which were exclusively present in flies from control and selected regimes, respectively. The unique DEGs identified in the selected flies may thus comprise genes that exhibit altered expression in response to adaptation to protein deprivation. We employed KEGG enrichment analyses to investigate the differential regulation of metabolic pathways related to reproduction or immune function. While we found downregulation in glycolysis, TCA cycle, pentose phosphate pathway, amino acid metabolism, and peroxisomal pathways in selected flies (**Figure 3C**), they also show lower activation of Imd and Toll pathways after infection.

To disentangle the response to infection as a function of selection for early-life fecundity, we next considered identifying DEGs with well-characterised functions relevant to reproduction and immunity. We identified 114 and 37 DEGs related to immune response and reproduction, respectively, in flies across all the infection and selection treatments. We divided the DEGs related to immune response into six broad categories, namely, (a) immune receptors, (b) immune regulators and (c) immune effectors driving inducible immunity, markers for (d) melanization and coagulation, (e) encapsulation and phagocytosis, and (f) ROS mediated defence (Westlake et al., 2024) (**Figure 3D**), which may evolve differently under pathogen selection (Seal et al, 2025). We have also identified DEGs that are known to be differentially regulated in response to infection, but their role in the immune response is not yet well-characterised (Westlake et al., 2024) (**Figure 3D**).

We further explored the associations of these DEGs from immune and reproductive gene categories as linearised metagene expression profiles using canonical variate analysis (CVA) to examine contrasting effects of infection treatments across selection regimes (**Figure 4**). We found a significant difference in post-infection gene expression profiles of all immune-related DEGs across regimes, with selected flies showing greater changes relative to their control (**Figure 4A, Table S17**). We also found pronounced divergence between sham-infected individuals from controls vs selected flies, suggesting potential background evolution due to selection (**Figure 4A**). Given this divergence in how infection affected total sets of immune genes across selection regimes, we subsequently analysed various functional immune categories separately to identify the relevant immune responses explaining the evolved phenotypic variations. Among the inducible immune components, control flies exhibited greater changes in gene expression profiles of immune receptors in response to infection (CV1 explaining 94.9% variance), primarily driven by Gram-negative bacteria-binding proteins (GNBPs) and peptidoglycan recognition proteins (PGRPs) (**Figure 4B, Table S18**). In contrast, selected flies showed greater alterations in the expression profile of regulatory genes (CV1 explaining 97.26% variance) (**Figure 4C, Table S19**), influenced by LUBEL, pirk, spirit, kappa B-Ras, relish, PGRP-SC2, helical factor and serine-type endopeptidase. Next, we compared expression profiles of effectors from inducible immune pathways using both CV1 and CV2, explaining >95% of variance (**Figure 4D–4E, Table S20**). CV1 reflected similar patterns of changes in gene expression patterns as found in receptors, with control flies showing greater divergence than selected flies, driven by Toll- and Imd-regulated AMPs such as cecropins, bomanin tailed-1, metchnikowin-like proteins, drosomycin-like proteins and defensin, immune-induced molecules such as IBIN (induced by infection) and edin, and putative AMP- and lysozyme-like molecules. In contrast, CV2 highlighted distinct sets of molecules that drove the divergence between control vs selected flies. For example, while the response of selected flies was driven by bomanin tailed-1, cecropin A1, IBIN, defensin, and lysozyme-like molecules, control flies showed changes driven by cecropin A2 & B, putative AMP-molecules, edin and drosomycin-like 4. We also compared the differential expression changes of individual driver genes within each immune category, revealing that selected flies typically showed lower activation than control flies (**Figure 3D**).

**Figure 4:**
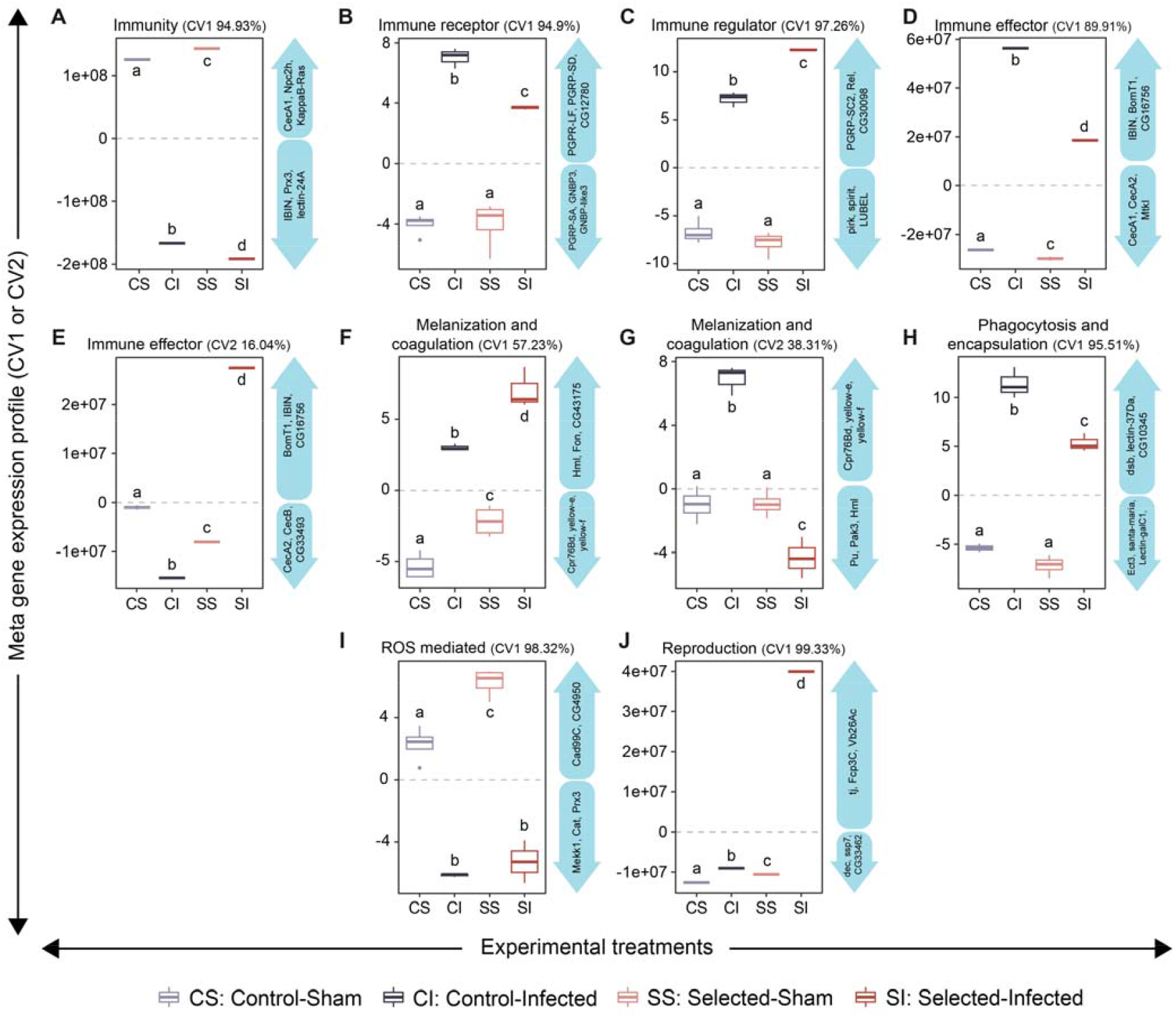
Canonical variate analyses (CVA) showing changes in gene expression profile of immune and reproductive genes as a function of infection and adaptation to chronic protein malnourishment. Cumulative gene expression profile based on canonical variate analysis (CVA) of differentially expressed genes, involved in (**A**) immune function, further categorized into: (**B**) receptors, (**C**) regulators, and (**D**) effectors driving inducible immune response, (**E**,**F**) melanization and coagulation, (**H**) phagocytosis and encapsulation, (**I**) ROS mediated responses. Additionally, Canonical variate analysis (CVA) performed on differentially expressed genes, involved in (**J**) reproduction. The change in expression profile is due to sham-infection or *Providencia rettgeri* infection in control vs selected regimes. The pairwise comparisons between different combinations of selection regimes and infection treatments are obtained from a post hoc test, where statistically significant differences between experimental treatment are indicated with different alphabets. The gene symbols of DEGs driving the changes in the metagene expression profiles are also mentioned within directed blue arrows at the right side of each panel.

We also included both CV1 and CV2 to explain the >95% variance of gene expression profile of melanisation and coagulation responses, driven by yellow-e, yellow-f, hemolectin, fondue, cuticular protein 76Bd. Although infection appeared to drive the main divergence in both selected and control flies along the CV1, there were also significant changes between their sham-infected treatments, suggesting interactions between the background variation and the effects of infection across selection regimes. However, CV2 proposed different patterns of contrast after infection across selection regimes, similar to those seen in inducible effectors (see above). While the response of selected flies was driven by punch, p21-activated kinase 3 (pak3) and hemolectin, control flies showed changes driven by yellow-e, yellow-f and cuticular protein 76Bd (**Figure 4F– 4G, Table S21**). Moreover, control flies exhibited greater changes in encapsulation and phagocytosis responses (CV1 explaining 95.5% variance), primarily driven by nimrodC1, Rab21, scavenger receptor ortholog CG10345, santa-maria, lectins, ectoderm-express 3, debris-buster (**Figure 4H, Table S22**). In contrast, selected flies showed greater divergence in ROS-related gene expression profile, driven by the negative regulator of ROS (nrROS) and cadherin 99C (**Figure 4I, Table S23**). We also found that, similar to inducible immune responses, several genes related to melanisation, coagulation, encapsulation, phagocytosis and ROS-mediated responses showed lower activation in selected flies as compared to their control counterparts (**Figure 3D**).

Finally, we also examined the changes in expression of reproduction-related DEGs. By and large, selected flies showed greater divergence in response to infection, including a significant background variation between sham-infected flies across regimes. The response was mainly driven by vitelline membrane protein 26Ac, traffic jam (tj), follicle cell protein 3C (fcp3c), putative serine-type endopeptidase CG33462 and short spindle 7 (ssp7) (**Figure 4J, Table S24**). Similar to inducible immunity, most of these reproduction-related genes were significantly more downregulated in selected flies than in controls after infection (**Figure 3D**).

## DISCUSSION

In this study, we investigated whether or to what extent the evolutionary history of chronic protein deprivation and strong selection for early-life reproduction drives variation in immunity and infection outcomes. While there was a clear trend for flies evolving under protein deprivation to show higher vulnerability to pathogenic bacteria, the level of trade-off varied considerably across sexes and mating status. For example, virgin females from the selected regime were more susceptible to *Providencia rettgeri* infection than their control counterparts. In contrast, effects on virgin males were weak and only dose-specific. Interestingly, following mating, individuals from both sexes from the selected regimes consistently exhibited increased susceptibility to infection, and the sex-specific effects faded away. We also note an interesting observation that immune strategies may differ widely across mating status. While assessed as virgins, selected females carried a higher pathogen burden, suggesting poor pathogen resistance and corroborating their increased post–infection mortality. However, no such difference was observed when the flies were mated, implying that the increased vulnerability of selected flies to pathogens may arise from a relative decline in their ability to withstand the pathogen load, i.e., reduced tolerance (Ayres and Schneider, 2012). The outcomes of reproduction-infection response trade-offs may thus depend on sex and mating status-specific variations in immune strategy.

A more pronounced and consistent decrease in post–infection survival among virgin females compared to males is expected, since selection for early-life fitness tends to be stronger in females within these fly populations (Dasgupta et al., 2022). This is because protein limitation is especially important for females, who need more protein for egg production (Mirth et al., 2019). On the other hand, although male reproductive effort is also protein-dependent, it is unlikely to be as intensive as in females, thus generating the sex difference in selection (Reddiex et al., 2013; Camus et al., 2018). Moreover, females from the selected regime in our experiments have been previously shown to have larger ovaries and a greater number of mature, post-vitellogenic oocytes, which may contribute to their higher early-life fecundity (Dasgupta et al., 2024). Although not empirically tested, they may also invest more in germline maintenance, as in many species, post–meiotic DNA damage repair mechanisms are non-functional in males (Olsen et al., 2005). All these, in turn, may lead to the adaptive reallocation of available protein reserves away from somatic maintenance (McKean and Nunney, 2008), precluding the ability to induce effective immune responses and prevent post– infection mortality, especially in females.

Why did the sexually dimorphic infection outcome disappear in mated evolved flies? Before answering this, it is essential to note that, as observed in various other taxa, mating alone appears to increase susceptibility to infection (Short and Lazzaro, 2010; Oku et al., 2019). Since mating triggers reproductive allocation by ramping up egg production in females (Ram and Wolfner, 2007) and stimulating seminal fluid and sperm production in males (Harvanek et al., 2017; Corbel et al., 2022), a more pronounced exhibition of immune trade–offs post–mating may manifest in both sexes. Literature on the effect of mating on reproductive investment appears to indicate the same, and importantly, in both sexes (Kraaijeveld and Godfray, 2008). Hence, it is not unreasonable to argue that a general mating-induced decline in overall post–infection survivorship may mask sex-specific differences in baseline immune investment and ability to survive infection. Here, the sex difference in immunity under virgin conditions may appear counterintuitive at first. Importantly, virgin condition represents the baseline, untriggered state. Therefore, our finding that the selection has a stronger impact on virgin immunity in females compared to males appears to reflect the evolution of the baseline resource allocation, which is more in line with the predictions from life-history and sexual selection theories.

What mechanisms might explain the observed trade-off between immunity and infection outcomes in selected flies? To answer this, we began by identifying significantly different gene expression profiles related to key immune, reproductive, and other metabolic pathways in selected flies compared to control individuals. A striking pattern emerged in meta gene expression profiles related to inducible immune components controlled by the IMD and Toll pathways, melanisation, phagocytosis, and ROS-mediated responses, which may have primarily driven the variation in infection response between the control and selected populations. While the selected flies may have failed to induce Toll- and IMD-responsive AMPs at levels necessary for effective immunity against bacterial protection, they also downregulated several genes associated with phagocytosis pathway, the melanisation pathway, and ROS-mediated defense more than what was observed in control flies, suggesting an overall lower immune competence. Similar decline in immune induction was observed in different insect species when they were reared on diets with qualitative or quantitative changes in dietary protein. For example, lysozyme-like antibacterial activity and PO activity were lower for *Spodoptera littoralis* caterpillars, reared on a low-quality zein-protein-supplemented diet (Lee et al., 2008). On the other hand, protein-deprived Bumble bees, *Bombus terrestris*, failed to upregulate *defensin* from the Toll pathway, *punch-1* from the melanisation pathway, and *jafrac* from the ROS-mediated pathway against parasitic infection caused by *Crithidia bombi* (Brunner et al., 2014).

Moreover, in the present study, the bacterial infection in selected flies drove a different set of DEGs related to inducible effectors (e.g., different antimicrobial peptides) and the melanisation response compared to their control counterparts. In control flies, gene expression of effector molecules was primarily driven by cecropin A2 & B, which could provide specific protection against the Gram-negative bacteria (Carboni et al., 2022). In selected flies, however, it was driven by antimicrobial peptides such as defensin, bomanin tailed-1, or IBIN. These peptides are also known for defending against Gram-positive bacteria, fungi, and parasitoid wasps (Nakajima et al., 2003; Westlake et al., 2022; Ebrahim et al., 2021), but their non-specificity in targeting Gram-negative bacterial pathogens can lead to suboptimal immune protection with increased cytotoxicity (Shit et al., 2022). Additionally, in selected flies, melanization response was primarily driven by genes, such as *punch*, which generates BH-4 (tetrahydrobiopterin) cofactor for phenylalanine hydroxylase that functions in the early stages of melanin production (Westlake et al., 2022; Chen et al., 2013); whereas in control flies, it was driven by *yellow e* and *yellow f* genes, which converts dopachrome to 5,6-Dihydroxyindole and are functional in the later stages of melanization pathway (Sugumaran and Barek, 2016). Thus, by controlling the expression of *punch* at a much earlier stage, selected flies could restrict energy utilised for the subsequent steps of the melanisation pathway and phenoloxidase response under protein-deprived conditions. This not only emphasises their inherent mechanistic differences in immune strategies and infection responses but may also underscore the molecular basis of why selected flies have weaker immunity than the control flies.

We also found downregulation in multiple metabolic pathways, including energy metabolism (Soto-Heredero et al., 2020) and biosynthetic pathways, such as amino acid (Kelly and Pearce, 2020), and peroxisome (Di Cara et al., 2018), in selected flies after infection, which has been shown in previous studies to be associated with response to pathogenic infections in various taxa. For example, in vertebrates, the metabolism of tryptophan and glutathione, which is composed of glutamate, glycine, and cysteine, is essential for T cell activation and the detoxification of ROS (Kelly and Pearce, 2020). Similarly, energy metabolism via glycolysis, the TCA cycle, or high lactate production is essential for providing energy to support inflammatory responses by macrophages and T lymphocytes (Soto-Heredero et al., 2020). This may have laid the metabolic basis for why selected flies were less competent in inducing immune responses and defending against infection; however, further experiments are needed to test these possibilities. Finally, there were significant alterations in the expression profile of reproduction-related genes in selected female flies after infection, relative to control flies, with notable downregulation of vitelline membrane proteins, serine racemase, and yolk proteins. Although not empirically tested, this may signify a greater reproductive cost of infection while adapting to chronic protein deprivation.

Another notable finding from our results is that the response to *P. rettgeri* infection in our fly populations under selection is not solely driven by Imd-responsive AMPs. This represents a significant deviation from previous studies using inbred fly lines, where Imd-responsive AMPs, such as Diptericin in males (Hanson et al., 2019) or Diptericin, Drosocin, and Attacins in females (Shit et al., 2022), have been shown to influence variation in infection outcomes critically. In fact, using CRISPR gene editing, these studies have even definitively demonstrated that the survival costs of *P. rettgeri* infection can be entirely reversed by restoring Imd-responsive AMPs against the genetic background of null mutants lacking all AMPs. However, our results provided additional insights. For instance, as expected, selected flies showed lower levels of Diptericin or Attacin expression following *P. rettgeri* infection, which could contribute to their increased susceptibility to this pathogen, but at the same time, these flies also displayed increased expression of Toll-responsive Defensin and Immune-induced molecule 18, both of which are effective against Gram-positive bacteria and fungi (Lemaitre et al., 1996; Anderson, 2000). Control flies also exhibited significant upregulation of Toll-responsive antimicrobial peptides (AMPs), including *Defensin* and *Cecropins*, implying that the exclusivity of Imd-responsive AMP-driven protection against *P. rettgeri* infection may not be a general phenomenon across populations and different genetic backgrounds.

Before we conclude, we would like to highlight that we measured bacterial load and gene expression only during the onset of mortality, representing the acute phase of infection. However, there can be significant temporal variations in gene expression patterns (Schlamp et al., 2021), which can result from changes in within-host bacterial density. Future studies should thus monitor the temporal dynamics of the bacterial load and its effects on immune induction and pathogen clearance to understand the observed post–infection survival patterns. Also, our findings were derived after one generation of relaxed selection pressure, which, while potentially indicating the effects of genetic adaptation, does not entirely rule out the possibility of non-genetic parental influences that can drive variations after several generations (Mondotte et al. 2020).

In summary, our results may be applicable to various wild species in their natural habitats (Carciofi and Saad, 2001), including humans in many impoverished nations (Semba, 2016), where access to adequate protein sources may be limited or intermittent, driving strong selection to rewire metabolic needs for growth and reproduction. Additionally, the possibility that an evolutionary background of chronic protein shortage compromises immune competence in extant generations, which can significantly impact the ability to tackle new pathogens, may prompt future epidemiological, immunological and health research to further confirm the repeatability of our results in other species within the context of disease ecology and emerging infections.

## Supporting information

Sarkar et al 2025_Supplementary information

## Availability of data and materials

Phenotypic data have been uploaded in Supplementary information 2. List of DEGs have been uploaded in Supplementary information 3. All the Raw RNA-seq data will be available in NCBI under Bioproject ID: PRJNA1282488.

## Authors’ contributions

Conceptualisation: IK, BN

Project administration: IK

Funding acquisition: IK

Resources: IK, BN

Supervision: IK

Methodology: SS, DNB, IK

Investigation: SS, SSe

Visualisation: SS, DNB

Validation: SS

Data curation: SS, DNB

Formal analysis: DNB, SS, IK

Writing – original draft: IK, SS, DNB

Writing – review & editing: IK, DNB, BN

## Acknowledgements

We thank Athulya Girish Kizhakke, Biswajit Shit, Muhammed Nirjas, and Selah Makinishi for their important feedback on the manuscript.

