## Supplementary material for "ADAPTATION AGAINST LONG-TERM PROTEIN DEPRIVATION TRADES OFF WITH IMMUNOCOMPETENCE AND THE ABILITY TO SURVIVE PATHOGENIC INFECTIONS": Sarkar et al 2025_Supplementary information

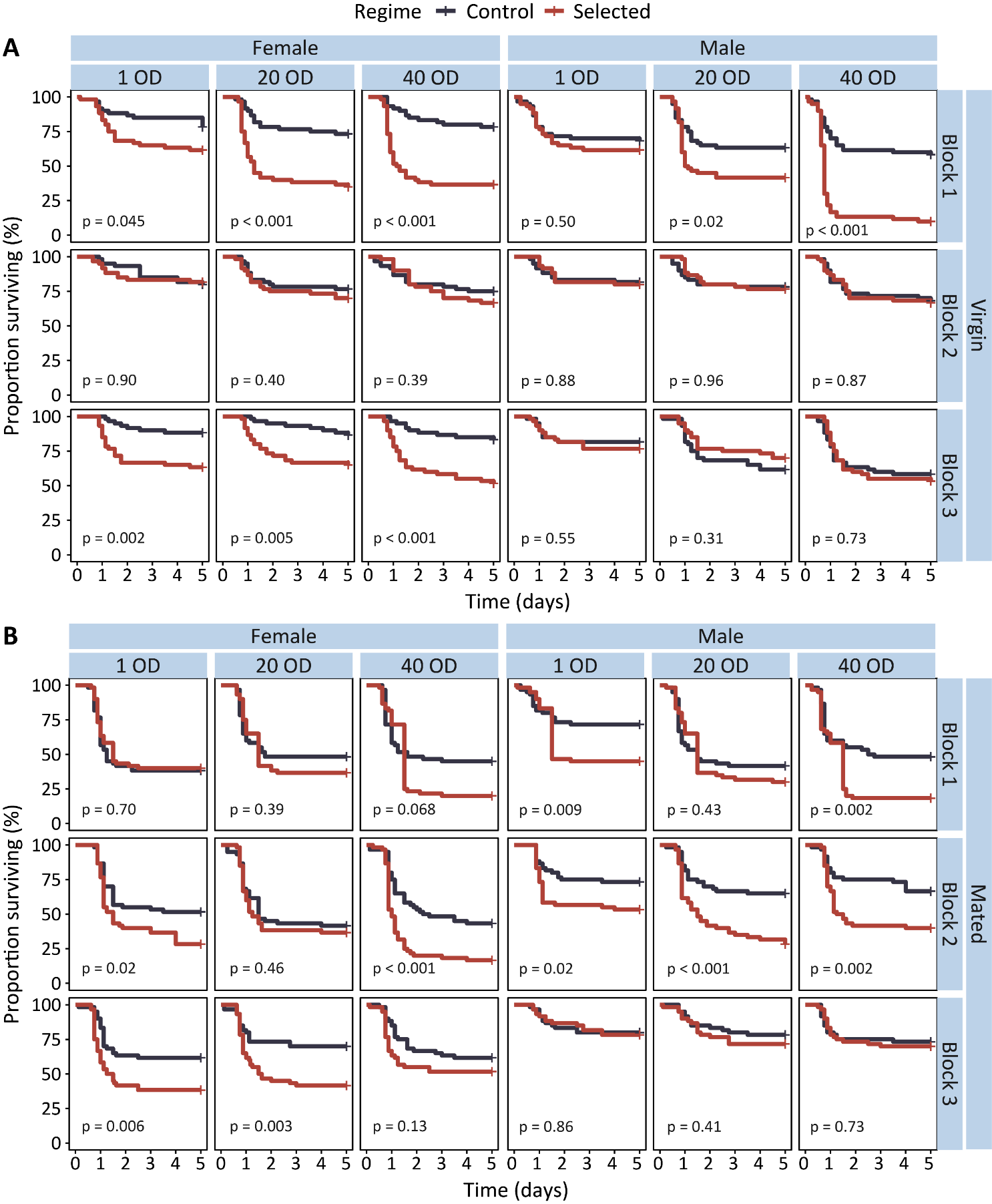
**Supplementary information**

**Figure S1.** Post-infection survival of replicate blocks of **(A)** virgin and **(B)** mated Drosophila melanogaster flies from control and selected regimes until 5 days post- *Providencia rettgeri* infection. The p values indicate the difference between control and selected regimes after *P. rettgeri* infection.

**TABLES**

**Table S1:** **Summary statistics of reduced model mixed-effect cox proportional hazard (coxme) analysis on post-infection survival of control and selected regimes.** The table shows the results of a reduced model coxme analysis, using infection dose (ID), selection regime (SR), mating status (MS) and sex (S) as fixed effects and vial identity nested within replicate populations as a random effect (Model: *Post-infection survival ~ Infection dose [ID]* *+* *Selection regime [SR] + Mating status [MS] + Sex [S] + ID×SR + ID×MS + ID×S + SR×MS + SR×S + MS×S + IS×MS×S + SR×MS×S + 1|* *Block / Vial*). Statistically significant p-values are highlighted in bold.

| Reduced  model | **Random factor** |  | sd | variance | |
| --- | --- | --- | --- | --- | --- |
|  | *Block/Vial* |  | 0.007 | < 0.001 | |
|  | *Block* |  | 0.333 | < 0.001 | |
|  | ***Tested effect*** |  | *Chisq* | *Df* | *p-value* |
|  | *Infection dose (ID)* |  | 463.116 | 3 | **< 0.001** |
|  | *Selection regime (SR)* |  | 115.502 | 1 | **< 0.001** |
|  | *Mating status (MS)* |  | 110.705 | 1 | **< 0.001** |
|  | *Sex (S)* |  | 6.521 | 1 | **0.011** |
|  | *ID ×* ***SR*** |  | 15.093 | 3 | **0.002** |
|  | *ID ×* *MS* |  | 17.614 | 3 | **< 0.001** |
|  | *ID × S* |  | 15.964 | 3 | **0.001** |
|  | *SR ×* *MS* |  | 2.537 | 1 | 0.111 |
|  | *SR ×* *S* |  | 1.435 | 1 | 0.231 |
|  | *MS × S* |  | 40.151 | 1 | **< 0.001** |
|  | *ID ×* *MS × S* |  | 10.157 | 3 | **0.017** |
|  | *SR ×* *MS ×* *S* |  | 9.064 | 1 | **0.003** |

**Table S2:** **Summary statistics of mixed effect cox proportional hazard (coxme) analysis on post-infection survival of virgin and mated flies from control and selected regimes.** The table shows the results of coxme analysis, separately performed for female and male flies, using infection dose (ID), selection regime (SR), and sex (S) as fixed effects and vial identity nested within replicate populations as a random effect (Model: Virgin- *Post-infection survival ~ Infection dose [ID]* *+* *Selection regime [SR] + Sex [S] + ID×SR + ID×S + SR×S + 1|* *Block / Vial*; Mated- *Post-infection survival ~ Infection dose [ID]* *+* *Selection regime [SR] + Sex [S] + ID×S + 1|* *Block / Vial*). Statistically significant p-values are highlighted in bold.

|  | |  | **Virgin** | | |  | **Mated** | | |
| --- | --- | --- | --- | --- | --- | --- | --- | --- | --- |
| Reduced  model | ***Random factor*** |  | *sd* | *variance* | |  | *sd* | *variance* | |
|  | *Block/Vial* |  | 0.006 | < 0.001 | |  | 0.012 | < 0.001 | |
|  | *Block* |  | 0.444 | < 0.001 | |  | 0.356 | 0.127 | |
|  | ***Tested effect*** |  | *Chisq* | *Df* | *p-value* |  | *Chisq* | *Df* | *p-value* |
|  | *Infection dose (ID)* |  | 210.693 | 3 | **< 0.001** |  | 278.689 | 3 | **< 0.001** |
|  | *Selection regime (SR)* |  | 65.920 | 1 | **< 0.001** |  | 56.994 | 1 | **< 0.001** |
|  | *Sex (S)* |  | 12.412 | 1 | **< 0.001** |  | 37.840 | 1 | **< 0.001** |
|  | *ID ×* *SR* |  | 14.265 | 3 | **0.002** |  | — | — | — |
|  | *ID × S* |  | 11.700 | 3 | **0.008** |  | 15.118 | 3 | **0.002** |
|  | *SR ×* *S* |  | 10.026 | 1 | **0.002** |  | — | — | — |

**Table S3:** **Summary statistics of mixed effect cox proportional hazard (coxme) analysis on post-infection survival of virgin female and male flies from control and selected regimes.** The table shows the results of coxme analysis, separately performed for virgin female and male flies, using infection dose (ID) and selection regime (SR) as fixed effects and vial identity nested within replicate populations as a random effect (Model: *Post-infection survival ~ Infection dose [ID]* *×* *Selection regime [SR] + 1|Block/Vial*). Statistically significant p-values are highlighted in bold.

| **Virgin** | | |  | **Female** | | | | | |  | **Male** | | | | | |
| --- | --- | --- | --- | --- | --- | --- | --- | --- | --- | --- | --- | --- | --- | --- | --- | --- |
| Full model | ***Random factor*** |  | | *sd* | | *variance* | | | |  | *sd* | | *variance* | | | |
|  | *Block/Vial* |  | | *0.020* | | < 0.001 | | | |  | *0.115* | | *0.013* | | | |
|  | *Block* |  | | *0.340* | | *0.116* | | | |  | *0.530* | | *0.281* | | | |
|  | ***Tested effect*** |  | | *Chisq* | | *Df* | | *p-value* | |  | *Chisq* | | *Df* | | *p-value* | |
|  | *Infection dose (ID)* |  | | 73.221 | | 3 | | **< 0.001** | |  | 146.667 | | 3 | | **< 0.001** | |
|  | *Selection Regime (SR)* |  | | 58.006 | | 1 | | **< 0.001** | |  | 17.246 | | 1 | | **< 0.001** | |
|  | *ID ×* ***SR*** |  | | 8.220 | | 3 | | **0.042** | |  | 8.316 | | 3 | | **0.040** | |
| Post hoc | *ID* |  | | *Estimate* | *SE* | | *Z ratio* | | *p-value* |  | *Estimate* | *SE* | | *Z ratio* | | *p-value* |
|  | *1 OD* |  | | -0.689 | 0.222 | | -3.109 | | **0.002** |  | -0.194 | 0.212 | | -0.914 | | 0.361 |
|  | *20 OD* |  | | -0.966 | 0.198 | | -4.877 | | **< 0.001** |  | -0.227 | 0.180 | | -1.267 | | 0.205 |
|  | *40 OD* |  | | -1.121 | 0.195 | | -5.751 | | **< 0.001** |  | -0.761 | 0.158 | | -4.819 | | **< 0.001** |

**Table S4:** **Summary statistics of mixed effect cox proportional hazard (coxme) analysis on post-infection survival of mated female and male flies from control and selected regimes.** The table shows the results of coxme analysis, separately performed for mated female and male flies, using infection dose and selection regime (SR) as fixed effects and vial identity nested within replicate populations as a random effect (Model: *Post-infection survival ~ Infection dose (ID)* *×* *Selection regime (SR) + 1|* *Block / Vial*). Statistically significant p-values are highlighted in bold.

| **Mated** | | |  | **Female** | | | | | |  | **Male** | | | | | |
| --- | --- | --- | --- | --- | --- | --- | --- | --- | --- | --- | --- | --- | --- | --- | --- | --- |
| Full model | ***Random factor*** |  | | *sd* | | *variance* | | | |  | *sd* | | *variance* | | | |
|  | *Block/Vial* |  | | *0.020* | | < 0.001 | | | |  | *0.020* | | < 0.001 | | | |
|  | *Block* |  | | *0.214* | | *0.046* | | | |  | *0.555* | | *0.308* | | | |
|  | ***Tested effect*** |  | | *Chisq* | | *Df* | | *p-value* | |  | *Chisq* | | *Df* | | *p-value* | |
|  | *Infection dose (ID)* |  | | 159.551 | | 3 | | **< 0.001** | |  | 136.43 | | 3 | | **< 0.001** | |
|  | *Selection Regime (SR)* |  | | 27.113 | | 1 | | **< 0.001** | |  | 32.64 | | 1 | | **< 0.001** | |
|  | *ID ×* ***SR*** |  | | 2.546 | | 3 | | 0.467 | |  | 2.70 | | 3 | | 0.441 | |
| Post hoc | *ID* |  | | *Estimate* | *SE* | | *Z ratio* | | *p-value* |  | *Estimate* | *SE* | | *Z ratio* | | *p-value* |
|  | *1 OD* |  | | -0.372 | 0.141 | | -2.636 | | **0.008** |  | -0.597 | 0.189 | | -3.154 | | **0.002** |
|  | *20 OD* |  | | -0.370 | 0.145 | | -2.557 | | **0.011** |  | -0.485 | 0.156 | | -3.107 | | **0.002** |
|  | *40 OD* |  | | -0.557 | 0.138 | | -4.035 | | **< 0.001** |  | -0.627 | 0.157 | | -3.987 | | **< 0.001** |

Table S5: Summary statistics of full model mixed-effect cox proportional hazard (coxme) analysis on post-infection survival of virgin female flies from control and selected regimes.

The table shows the results of coxme analyses on post-infection survival data of virgin female flies of different replicate blocks using infection dose (ID), selection regime (SR) as fixed effects and vial identity as a random effect (Model: *Post-infection survival ~ Infection dose [ID] ×* *Selection regime [SR] + 1|Vial*). Statistically significant p-values are highlighted in bold.

| **Virgin**  **female** | Block 1 | Full model | **Tested effect** |  | Chisq | Df | | p-value |
| --- | --- | --- | --- | --- | --- | --- | --- | --- |
|  |  |  | *Infection Dose (ID)* |  | 39.923 | 3 | | **< 0.001** |
|  |  |  | *Selection regime (SR)* |  | 38.743 | 1 | | **< 0.001** |
|  |  |  | *ID × SR* |  | 6.657 | 3 | | 0.084 |
|  |  | Post hoc  comparisons | *ID* |  | *estimate* | *SE* | *Z ratio* | *p-value* |
|  |  |  | *1 OD* |  | -0.695 | 0.347 | -2.001 | **0.045** |
|  |  |  | *20 OD* |  | -1.287 | 0.297 | -4.327 | **< 0.001** |
|  |  |  | *40 OD* |  | -1.541 | 0.322 | -4.785 | **< 0.001** |
|  | *Bl*ock 2 | *Full model* | ***Tested effect*** |  | *Chisq* | *Df* | | *p-value* |
|  |  |  | *Infection Dose (ID)* |  | 18.034 | 3 | | **< 0.001** |
|  |  |  | *Selection regime (SR)* |  | 0.865 | 1 | | 0.352 |
|  |  |  | *ID × SR* |  | 0.619 | 3 | | 0.892 |
|  |  | Post hoc  comparisons | *ID* |  | *estimate* | *SE* | *Z ratio* | *p-value* |
|  |  |  | *1 OD* |  | 0.051 | 0.417 | 0.122 | 0.903 |
|  |  |  | *20 OD* |  | -0.303 | 0.356 | -0.850 | 0.395 |
|  |  |  | *40 OD* |  | -0.295 | 0.342 | -0.864 | 0.387 |
|  | *Bl*ock 3 | *Full model* | ***Tested effect*** |  | *Chisq* | *Df* | | *p-value* |
|  |  |  | *Infection Dose (ID)* |  | 17.750 | 3 | | **< 0.001** |
|  |  |  | *Selection regime (SR)* |  | 27.259 | 1 | | **< 0.001** |
|  |  |  | *ID × SR* |  | 3.842 | 3 | | 0.279 |
|  |  | Post hoc  comparisons | *ID* |  | *estimate* | *SE* | *Z ratio* | *p-value* |
|  |  |  | *1 OD* |  | -1.36 | 0.434 | -3.130 | **0.002** |
|  |  |  | *20 OD* |  | -1.16 | 0.416 | -2.783 | **0.005** |
|  |  |  | *40 OD* |  | -1.35 | 0.367 | -3.688 | **< 0.001** |

Table S6: Summary statistics of full model mixed-effect cox proportional hazard (coxme) analysis on post-infection survival of virgin male flies from control and selected regimes.

The table shows the results of coxme analyses on post-infection survival data of virgin male flies of different replicate blocks using infection dose (ID), selection regime (SR) as fixed effects and vial identity as a random effect (Model: *Post-infection survival ~ Infection dose [ID] ×* *Selection regime [SR] + 1|Vial*). Statistically significant p-values are highlighted in bold.

| **Virgin**  **male** | Block 1 | Full model | **Tested effect** |  | Chisq | Df | | p-value |
| --- | --- | --- | --- | --- | --- | --- | --- | --- |
|  |  |  | *Infection Dose (ID)* |  | 87.973 | 3 | | **< 0.001** |
|  |  |  | *Selection regime (SR)* |  | 30.611 | 1 | | **< 0.001** |
|  |  |  | *ID × SR* |  | 14.687 | 3 | | **0.002** |
|  |  | Post hoc  comparisons | *ID* |  | *estimate* | *SE* | *Z ratio* | *p-value* |
|  |  |  | *1 OD* |  | -0.217 | 0.310 | -0.701 | 0.483 |
|  |  |  | *20 OD* |  | -0.639 | 0.273 | -2.343 | **0.019** |
|  |  |  | *40 OD* |  | -1.549 | 0.246 | -6.293 | **< 0.001** |
|  | *Bl*ock 2 | *Full model* | ***Tested effect*** |  | *Chisq* | *Df* | | *p-value* |
|  |  |  | *Infection Dose (ID)* |  | 20.682 | 3 | | **< 0.001** |
|  |  |  | *Selection regime (SR)* |  | 0.045 | 1 | | 0.832 |
|  |  |  | *ID × SR* |  | 0.010 | 3 | | 0.9997 |
|  |  | Post hoc  comparisons | *ID* |  | *estimate* | *SE* | *Z ratio* | *p-value* |
|  |  |  | *1 OD* |  | -0.065 | 0.417 | -0.156 | 0.876 |
|  |  |  | *20 OD* |  | -0.028 | 0.385 | -0.072 | 0.943 |
|  |  |  | *40 OD* |  | -0.051 | 0.320 | -0.159 | 0.873 |
|  | *Bl*ock 3 | *Full model* | ***Tested effect*** |  | *Chisq* | *Df* | | *p-value* |
|  |  |  | *Infection Dose (ID)* |  | 40.296 | 3 | | **< 0.001** |
|  |  |  | *Selection regime (SR)* |  | 0.006 | 1 | | 0.940 |
|  |  |  | *ID × SR* |  | 1.572 | 3 | | 0.666 |
|  |  | Post hoc  comparisons | *ID* |  | *estimate* | *SE* | *Z ratio* | *p-value* |
|  |  |  | *1 OD* |  | -0.249 | 0.403 | -0.618 | 0.537 |
|  |  |  | *20 OD* |  | 0.325 | 0.315 | 1.031 | 0.302 |
|  |  |  | *40 OD* |  | -0.100 | 0.276 | -0.364 | 0.716 |

Table S7: Summary statistics of full model mixed-effect cox proportional hazard (coxme) analysis on post-infection survival of mated female flies from control and selected regimes.

| **Mated**  **female** | Block 1 | Full model | **Tested effect** |  | Chisq | Df | | p-value |
| --- | --- | --- | --- | --- | --- | --- | --- | --- |
|  |  |  | *Infection Dose (ID)* |  | 60.469 | 3 | | **< 0.001** |
|  |  |  | *Selection regime (SR)* |  | 1.71 | 1 | | 0.191 |
|  |  |  | *ID × SR* |  | 2.481 | 3 | | 0.479 |
|  |  | Post hoc  comparisons | *ID* |  | *estimate* | *SE* | *Z ratio* | *p-value* |
|  |  |  | *1 OD* |  | 0.090 | 0.234 | 0.384 | 0.701 |
|  |  |  | *20 OD* |  | -0.208 | 0.242 | -0.857 | 0.392 |
|  |  |  | *40 OD* |  | -0.414 | 0.227 | -1.822 | 0.068 |
|  | *Bl*ock 2 | *Full model* | ***Tested effect*** |  | *Chisq* | *Df* | | *p-value* |
|  |  |  | *Infection Dose (ID)* |  | 68.130 | 3 | | **< 0.001** |
|  |  |  | *Selection regime (SR)* |  | 15.102 | 1 | | **< 0.001** |
|  |  |  | *ID × SR* |  | 5.072 | 3 | | 0.167 |
|  |  | Post hoc  comparisons | *ID* |  | *estimate* | *SE* | *Z ratio* | *p-value* |
|  |  |  | *1 OD* |  | -0.550 | 0.241 | -2.287 | **0.022** |
|  |  |  | *20 OD* |  | -0.172 | 0.234 | -0.734 | 0.463 |
|  |  |  | *40 OD* |  | -0.849 | 0.223 | -3.805 | **< 0.001** |
|  | *Bl*ock 3 | *Full model* | ***Tested effect*** |  | *Chisq* | *Df* | | *p-value* |
|  |  |  | *Infection Dose (ID)* |  | 38.879 | 3 | | **< 0.001** |
|  |  |  | *Selection regime (SR)* |  | 16.323 | 1 | | **< 0.001** |
|  |  |  | *ID × SR* |  | 2.383 | 3 | | 0.497 |
|  |  | Post hoc  comparisons | *ID* |  | *estimate* | *SE* | *Z ratio* | *p-value* |
|  |  |  | *1 OD* |  | -0.728 | 0.266 | -2.738 | **0.006** |
|  |  |  | *20 OD* |  | -0.871 | 0.290 | -2.999 | **0.003** |
|  |  |  | *40 OD* |  | -0.419 | 0.279 | -1.502 | 0.133 |

The table shows the results of coxme analyses on post-infection survival data of mated female flies of different replicate blocks using infection dose (ID), selection regime (SR) as fixed effects and vial identity as a random effect (Model: *Post-infection survival ~ Infection dose [ID] ×* *Selection regime [SR] + 1|Vial*). Statistically significant p-values are highlighted in bold.

Table S8: Summary statistics of full model mixed-effect cox proportional hazard (coxme) analysis on post-infection survival of mated male flies from control and selected regimes.

The table shows the results of coxme analyses on post-infection survival data of mated male flies of different replicate blocks using infection dose (ID), selection regime (SR) as fixed effects and vial identity as a random effect (Model: *Post-infection survival ~ Infection dose [ID] ×* *Selection regime [SR] + 1|Vial*). Statistically significant p-values are highlighted in bold.

| **Mated**  **male** | Block 1 | Full model | **Tested effect** |  | Chisq | Df | | p-value |
| --- | --- | --- | --- | --- | --- | --- | --- | --- |
|  |  |  | *Infection Dose (ID)* |  | 75.186 | 3 | | **< 0.001** |
|  |  |  | *Selection regime (SR)* |  | 12.830 | 1 | | **< 0.001** |
|  |  |  | *ID × SR* |  | 4.378 | 3 | | 0.223 |
|  |  | Post hoc  comparisons | *ID* |  | *estimate* | *SE* | *Z ratio* | *p-value* |
|  |  |  | *1 OD* |  | -0.779 | 0.299 | -2.607 | **0.009** |
|  |  |  | *20 OD* |  | -0.182 | 0.229 | -0.796 | 0.426 |
|  |  |  | *40 OD* |  | -0.725 | 0.231 | -3.144 | **0.002** |
|  | *Bl*ock 2 | *Full model* | ***Tested effect*** |  | *Chisq* | *Df* | | *p-value* |
|  |  |  | *Infection Dose (ID)* |  | 44.884 | 3 | | **< 0.001** |
|  |  |  | *Selection regime (SR)* |  | 26.795 | 1 | | **< 0.001** |
|  |  |  | *ID × SR* |  | 2.348 | 3 | | 0.503 |
|  |  | Post hoc  comparisons | *ID* |  | *estimate* | *SE* | *Z ratio* | *p-value* |
|  |  |  | *1 OD* |  | -0.722 | 0.314 | -2.302 | **0.021** |
|  |  |  | *20 OD* |  | -1.019 | 0.267 | -3.820 | **< 0.001** |
|  |  |  | *40 OD* |  | -0.855 | 0.279 | -3.061 | **0.002** |
|  | *Bl*ock 3 | *Full model* | ***Tested effect*** |  | *Chisq* | *Df* | | *p-value* |
|  |  |  | *Infection Dose (ID)* |  | 16.532 | 3 | | **< 0.001** |
|  |  |  | *Selection regime (SR)* |  | 0.550 | 1 | | 0.458 |
|  |  |  | *ID × SR* |  | 0.278 | 3 | | 0.964 |
|  |  | Post hoc  comparisons | *ID* |  | *estimate* | *SE* | *Z ratio* | *p-value* |
|  |  |  | *1 OD* |  | -0.072 | 0.400 | -0.180 | 0.857 |
|  |  |  | *20 OD* |  | -0.304 | 0.368 | -0.825 | 0.409 |
|  |  |  | *40 OD* |  | -0.117 | 0.344 | -0.339 | 0.735 |

**Table S9:** **Summary statistics of a reduced generalized linear mixed-effects model (GLMM) analysis on hazard ratios of selected vs control regimes.** The table shows the results of GLMM analysis, on hazard ratios (calculated by comparing selected flies to control flies post-infection), with infection dose (ID), mating status (MS) and sex (S) as fixed effects and vial identity nested within the replicate populations as a random effect (Model: *Hazard ratio ~ Mating status [MS] + Sex [S] + MS×S* *+ 1| Block /Vial, family=negative binomial*). Statistically significant p-values have been highlighted in bold.

| Reduced model | **Random factor** |  | sd | variance | |
| --- | --- | --- | --- | --- | --- |
|  | *Block/Vial* |  | < 0.001 | < 0.001 | |
|  | *Block* |  | < 0.001 | < 0.001 | |
|  | ***Tested effect*** |  | *Chisq* | *Df* | *p-value* |
|  | *Mating status (MS)* |  | 3.361 | 1 | 0.067 |
|  | *Sex (S)* |  | 4.366 | 1 | **0.037** |
|  | *MS × S* |  | 18.180 | 1 | **< 0.001** |

**Table S10:** **Summary statistics of a generalized linear mixed-effects model (GLMM) analysis on hazard ratios of selected vs control regimes at individual doses.** The table shows the results of a generalized linear mixed-effects model on hazard ratios (calculated by comparing selected flies to control flies post-infection), calculated at individual doses, with mating status (MS) and sex (S) as fixed effects and vial identity nested within the replicate populations as a random effect (Model: *Hazard ratio ~ Mating status (MS) × Sex [S]* *+ 1| Block /Vial, family=negative binomial*). Statistically significant p-values have been highlighted in bold.

| 1 OD | Full model | **Tested effect** |  | Chisq | Df | | p-value |
| --- | --- | --- | --- | --- | --- | --- | --- |
|  |  | *Mating status (MS)* |  | 0.024 | 1 | | 0.877 |
|  |  | *Sex (S)* |  | 0.219 | 1 | | 0.640 |
|  |  | *MS × S* |  | 5.774 | 1 | | **0.016** |
|  | Post hoc | *Sex* |  | *estimate* | *SE* | *Z ratio* | *p-value* |
|  |  | *Female* |  | 0.566 | 0.318 | 1.783 | 0.075 |
|  |  | *Male* |  | -0.537 | 0.330 | -1.628 | 0.104 |
| 20 OD | *Full model* | ***Tested effect*** |  | *Chisq* | *Df* | | *p-value* |
|  |  | *Mating status (MS)* |  | 2.405 | 1 | | 0.121 |
|  |  | *Sex (S)* |  | 5.042 | 1 | | **0.025** |
|  |  | *MS × S* |  | 9.167 | 1 | | **0.002** |
|  | Post hoc | *Sex* |  | *estimate* | *SE* | *Z ratio* | *p-value* |
|  |  | *Female* |  | 0.889 | 0.282 | 3.153 | **0.002** |
|  |  | *Male* |  | -0.414 | 0.325 | -1.272 | 0.203 |
| 40 OD | *Full model* | ***Tested effect*** |  | *Chisq* | *Df* | | *p-value* |
|  |  | *Mating status (MS)* |  | 2.625 | 1 | | 0.105 |
|  |  | *Sex (S)* |  | 1.251 | 1 | | 0.263 |
|  |  | *MS × S* |  | 4.006 | 1 | | **0.045** |
|  | Post hoc | *Sex* |  | *estimate* | *SE* | *Z ratio* | *p-value* |
|  |  | *Female* |  | 0.682 | 0.267 | 2.555 | **0.011** |
|  |  | *Male* |  | -0.088 | 0.278 | -0.318 | 0.751 |

**Table S11: Summary statistics of a reduced model generalized linear mixed-effects (GLMM) analysis on bacterial load of control and selected regimes.** The table shows the results of a reduced GLMM analysis on bacterial load, using infection dose (ID), selection regime (SR), mating status (MS) and sex (S) as fixed effects and vial identity nested within replicate populations as a random effect (Model: *Log bacterial load~ Infection dose [ID]* ***+*** *Selection regime [SR]* ***+*** *Mating status [MS]* ***+ Sex [S] + MS****×****S* *+ 1|Block/Vial****, family=negative binomial*). Statistically significant p-values have been highlighted in bold.

| Reduced model | **Random factor** |  | sd | variance | |
| --- | --- | --- | --- | --- | --- |
|  | *Block/Vial* |  | < 0.001 | < 0.001 | |
|  | *Block* |  | < 0.001 | < 0.001 | |
|  | ***Tested effect*** |  | *Chisq* | *Df* | *p-value* |
|  | *Infection dose (ID)* |  | 75.861 | 2 | **< 0.001** |
|  | *Selection regime (SR)* |  | 7.628 | 1 | **0.006** |
|  | *Mating status (MS)* |  | 10.138 | 1 | **0.001** |
|  | *Sex (S)* |  | 1.468 | 1 | 0.226 |
|  | *MS × S* |  | 3.691 | 1 | 0.055 |

**Table S12:** **Summary statistics of a reduced model generalized linear mixed-effects (GLMM) analysis on bacterial load of virgin and mated flies from control and selected regimes.** The table shows the results of a reduced GLMM analysis on bacterial load of virgin and mated flies using infection dose (ID), selection regime (SR), and sex (S) as fixed effects and vial identity nested within replicate populations as a random effect (Model: Virgin- *Log bacterial load ~ Infection dose [ID]* *+* *Selection regime [SR] + Sex [S] + ID×SR + 1|* *Block / Vial*; Mated- *Log bacterial load ~ Infection dose [ID]* *+ Sex [S] + 1|* *Block / Vial*). Statistically significant p-values are highlighted in bold.

|  | |  | **Virgin** | | |  | **Mated** | | |
| --- | --- | --- | --- | --- | --- | --- | --- | --- | --- |
| Reduced  model | ***Random factor*** |  | *sd* | *variance* | |  | *sd* | *variance* | |
|  | *Block/Vial* |  | < 0.001 | < 0.001 | |  | < 0.001 | < 0.001 | |
|  | *Block* |  | < 0.001 | < 0.001 | |  | 0.041 | 0.002 | |
|  | ***Tested effect*** |  | *Chisq* | *Df* | *p-value* |  | *Chisq* | *Df* | *p-value* |
|  | *Infection dose (ID)* |  | 60.233 | 2 | **< 0.001** |  | 23.231 | 2 | **< 0.001** |
|  | *Selection regime (SR)* |  | 6.980 | 1 | **0.008** |  | — | — | — |
|  | *Sex (S)* |  | — | — | — |  | 4.926 | 1 | **0.026** |
|  | *ID ×* *SR* |  | 6.244 | 2 | **0.044** |  | — | — | — |

**Table S13:** **Summary statistics of generalized linear mixed-effects (GLMM) analysis on bacterial load of virgin female and male flies from control and selected regimes.** The table shows the results of a GLMM analysis on bacterial load of virgin female and male flies using infection dose, and selection regime as fixed effects and vial identity nested within replicate populations as a random effect (Model: *Post-infection survival ~ Infection dose (ID)* *×* *Selection regime (SR) + 1|* *Block / Vial*). Statistically significant p-values are highlighted in bold.

| **Virgin** | | |  | **Female** | | | | | |  | **Male** | | | | | |
| --- | --- | --- | --- | --- | --- | --- | --- | --- | --- | --- | --- | --- | --- | --- | --- | --- |
| Full model | ***Random factor*** |  | | *sd* | | *variance* | | | |  | *sd* | | *variance* | | | |
|  | *Block/Vial* |  | | < 0.001 | | < 0.001 | | | |  | < 0.001 | | < 0.001 | | | |
|  | *Block* |  | | < 0.001 | | < 0.001 | | | |  | 0.070 | | 0.005 | | | |
|  | ***Tested effect*** |  | | *Chisq* | | *Df* | | *p-value* | |  | *Chisq* | | *Df* | | *p-value* | |
|  | *Infection dose (ID)* |  | | 32.568 | | 2 | | **< 0.001** | |  | 31.347 | | 2 | | **< 0.001** | |
|  | *Selection Regime (SR)* |  | | 8.150 | | 1 | | **0.004** | |  | 0.721 | | 1 | | 0.396 | |
|  | *ID ×* ***SR*** |  | | 3.657 | | 2 | | 0.161 | |  | 2.726 | | 2 | | 0.256 | |
| Post hoc | *ID* |  | | *Estimate* | *SE* | | *Z ratio* | | *p-value* |  | *Estimate* | *SE* | | *Z ratio* | | *p-value* |
|  | *1 OD* |  | | -0.319 | 0.104 | | -3.070 | | **0.002** |  | -0.172 | 0.101 | | -1.705 | | 0.088 |
|  | *20 OD* |  | | -0.067 | 0.088 | | -0.770 | | 0.441 |  | 0.049 | 0.091 | | 0.536 | | 0.592 |
|  | *40 OD* |  | | -0.128 | 0.091 | | -1.415 | | 0.157 |  | -0.044 | 0.085 | | -0.513 | | 0.608 |

**Table S14:** **Summary statistics of generalized linear mixed-effects (GLMM) analysis on bacterial load of mated female and male flies from control and selected regimes.** The table shows the results of a GLMM analysis on bacterial load mated female and male flies using infection dose, and selection regime as fixed effects and vial identity nested within replicate populations as a random effect (Model: *Post-infection survival ~ Infection dose (ID)* *×* *Selection regime (SR) + 1|* *Block / Vial*). Statistically significant p-values are highlighted in bold.

| **Mated** | | |  | **Female** | | | | | |  | **Male** | | | | | |
| --- | --- | --- | --- | --- | --- | --- | --- | --- | --- | --- | --- | --- | --- | --- | --- | --- |
| Full model | ***Random factor*** |  | | *sd* | | *variance* | | | |  | *sd* | | *variance* | | | |
|  | *Block/Vial* |  | | < 0.001 | | < 0.001 | | | |  | < 0.001 | | < 0.001 | | | |
|  | *Block* |  | | < 0.001 | | < 0.001 | | | |  | 0.059 | | 0.003 | | | |
|  | ***Tested effect*** |  | | *Chisq* | | *Df* | | *p-value* | |  | *Chisq* | | *Df* | | *p-value* | |
|  | *Infection dose (ID)* |  | | 9.831 | | 2 | | **0.007** | |  | 14.14 | | 2 | | **< 0.001** | |
|  | *Selection Regime (SR)* |  | | 0.674 | | 1 | | 0.412 | |  | 1.13 | | 1 | | 0.287 | |
|  | *ID ×* ***SR*** |  | | 1.775 | | 2 | | 0.412 | |  | 2.20 | | 2 | | 0.332 | |
| Post hoc | *ID* |  | | *Estimate* | *SE* | | *Z ratio* | | *p-value* |  | *Estimate* | *SE* | | *Z ratio* | | *p-value* |
|  | *1 OD* |  | | -0.058 | 0.093 | | -0.628 | | 0.530 |  | -0.005 | 0.092 | | -0.064 | | 0.949 |
|  | *20 OD* |  | | 0.046 | 0.087 | | 0.532 | | 0.595 |  | -0.161 | 0.088 | | -1.825 | | 0.068 |
|  | *40 OD* |  | | -0.114 | 0.085 | | -1.342 | | 0.180 |  | 0.004 | 0.088 | | 0.049 | | 0.961 |

Table S15: Summary statistics of generalized linear model (GLM) analysis on infection tolerance of virgin female and male flies from control and selected regimes.

The table shows the results of a GLM fitted for fitted separately for virgin female and male flies, using log-transformed hazard function as a response variable, pseudo-log-transformed average bacterial load (BL) as a covariate and selection regime (SR) as a fixed effect (Model: *Log hazard ratio~ Log average bacterial load (BL)* ***×*** *Selection regime (SR), family=negative binomial*). A significant interaction between bacterial load and selection regime indicates variation in infection tolerance, highlighted in bold.

| **Virgin** |  | **Female** | | |  | **Male** | | |
| --- | --- | --- | --- | --- | --- | --- | --- | --- |
| **Tested effect** |  | *Chisq* | *Df* | *p-value* |  | *Chisq* | *Df* | *p-value* |
| Log Avg bacterial load (BL) |  | 2.779 | 1 | 0.095 |  | 2.534 | 1 | 0.111 |
| Selection regime (SR) |  | 23.577 | 1 | < 0.001 |  | 3.906 | 1 | 0.048 |
| BL **× SR** |  | 3.270 | 1 | 0.071 |  | 7.419 | 1 | **0.006** |

Table S16: Summary statistics of generalized linear model (GLM) analysis on infection tolerance of mated female and male flies from control and selected regimes.

The table shows the results of a GLM fitted for fitted separately for mated female and male flies, using log-transformed hazard function (HR) as a response variable, pseudo-log-transformed average bacterial load (BL) as a covariate and selection regime (SR) as a fixed effect (Model: *Log hazard ratio~ Log average bacterial load* **×** *Selection regime, family=negative binomial*). A significant interaction between bacterial load and selection regime indicates variation in infection tolerance, highlighted in bold.

| **Mated** |  | **Female** | | |  | **Male** | | |
| --- | --- | --- | --- | --- | --- | --- | --- | --- |
| **Tested effect** |  | *Chisq* | *Df* | *p-value* |  | *Chisq* | *Df* | *p-value* |
| Log Avg bacterial load (BL) |  | 1.020 | 1 | 0.313 |  | 21.808 | 1 | < 0.001 |
| Selection regime (SR) |  | 24.906 | 1 | < 0.001 |  | 11.561 | 1 | < 0.001 |
| BL **× SR** |  | 1.881 | 1 | 0.170 |  | 1.113 | 1 | 0.291 |

**Table S17: Post hoc test statistics from CVA of immune genes.** This table shows differences between gene expression profiles of immune genes in flies from control and selected regimes post-infection.

| **Immunity**  **CV1** | **Experimental treatments** |  | diff | lower | upper | p-adj |
| --- | --- | --- | --- | --- | --- | --- |
|  | *Control sham-Control infected* |  | 292197930 | 292197930 | 292197930 | **< 0.001** |
|  | *Selected infected-Control infected* |  | -24697797 | -24697797 | -24697797 | **< 0.001** |
|  | *Selected sham-Control infected* |  | 309916088 | 309916088 | 309916088 | **< 0.001** |
|  | *Selected infected-Control sham* |  | -316895727 | -316895727 | -316895727 | **< 0.001** |
|  | *Selected sham- Control sham* |  | 17718158 | 17718158 | 17718158 | **< 0.001** |
|  | *Selected sham- Selected infected* |  | 334613885 | 334613885 | 334613885 | **< 0.001** |

**Table S18: Post hoc test statistics from CVA of immune-receptor genes.** This table shows differences between gene expression profiles of immune-receptor genes in flies from control and selected regimes post-infection.

| **Immune**  **receptor**  **CV1** | **Experimental treatments** |  | diff | lower | upper | p-adj |
| --- | --- | --- | --- | --- | --- | --- |
|  | *Control sham-Control infected* |  | -11.057149 | -13.393771 | -8.7205279 | **< 0.001** |
|  | *Selected infected-Control infected* |  | -3.336638 | -5.834591 | -0.8386843 | **0.009** |
|  | *Selected sham-Control infected* |  | -11.024232 | -13.360854 | -8.6876109 | **< 0.001** |
|  | *Selected infected-Control sham* |  | 7.720512 | 5.383890 | 10.0571331 | **< 0.001** |
|  | *Selected sham- Control sham* |  | 0.032917 | -2.130374 | 2.1962081 | 0.999 |
|  | *Selected sham- Selected infected* |  | -7.687595 | -10.024216 | -5.3509733 | **< 0.001** |

**Table S19: Post hoc test statistics from CVA of immune-regulator genes.** This table shows differences between gene expression profiles of immune-regulator genes in flies from control and selected regimes post-infection.

| **Immune**  **regulator**  **CV1** | **Experimental treatments** |  | diff | lower | upper | p-adj |
| --- | --- | --- | --- | --- | --- | --- |
|  | *Control sham-Control infected* |  | -13.868353 | -16.204974 | -11.531732 | **< 0.001** |
|  | *Selected infected-Control infected* |  | 5.166679 | 2.668725 | 7.664632 | **< 0.001** |
|  | *Selected sham-Control infected* |  | -15.006684 | -17.343306 | -12.670063 | **< 0.001** |
|  | *Selected infected-Control sham* |  | 19.035032 | 16.698410 | 21.371653 | **< 0.001** |
|  | *Selected sham- Control sham* |  | -1.138331 | -3.301622 | 1.024960 | 0.416 |
|  | *Selected sham- Selected infected* |  | -20.173363 | -22.509984 | -17.836742 | **< 0.001** |

**Table S20: Post hoc test statistics from CVA of immune-effector genes.** This table shows differences between gene expression profiles of immune-effector genes in flies from control and selected regimes post-infection.

| **Immune**  e**ffectors**  **CV1** | **Experimental treatments** |  | diff | lower | upper | p-adj |
| --- | --- | --- | --- | --- | --- | --- |
|  | *Control sham-Control infected* |  | -82606104 | -82606104 | -82606104 | **< 0.001** |
|  | *Selected infected-Control infected* |  | -37768561 | -37768561 | -37768561 | **< 0.001** |
|  | *Selected sham-Control infected* |  | -86222652 | -86222652 | -86222652 | **< 0.001** |
|  | *Selected infected-Control sham* |  | 44837544 | 44837544 | 44837544 | **< 0.001** |
|  | *Selected sham- Control sham* |  | -3616548 | -3616548 | -3616548 | **< 0.001** |
|  | *Selected sham- Selected infected* |  | -48454091 | -48454091 | -48454091 | **< 0.001** |
| **Immune**  **effectors**  **CV2** | *Control sham-Control infected* |  | 14365808 | 14365808 | 14365808 | **< 0.001** |
|  | *Selected infected-Control infected* |  | 42788332 | 42788332 | 42788332 | **< 0.001** |
|  | *Selected sham-Control infected* |  | 7298952 | 7298952 | 7298952 | **< 0.001** |
|  | *Selected infected-Control sham* |  | 28422525 | 28422525 | 28422525 | **< 0.001** |
|  | *Selected sham- Control sham* |  | -7066856 | -7066856 | -7066856 | **< 0.001** |
|  | *Selected sham- Selected infected* |  | -35489380 | -35489380 | -35489380 | **< 0.001** |

**Table S21: Post hoc test statistics from CVA of** **melanization and coagulation related immune genes.** This table shows differences between gene expression profiles of melanization and coagulation related immune genes in flies from control and selected regimes post-infection.

| **Melanization and**  **coagulation**  **CV1** | **Experimental treatments** |  | diff | lower | upper | p-adj |
| --- | --- | --- | --- | --- | --- | --- |
|  | *Control sham-Control infected* |  | -8.371379 | -10.708000 | -6.034757 | **< 0.001** |
|  | *Selected infected-Control infected* |  | 4.044724 | 1.546770 | 6.542677 | **0.002** |
|  | *Selected sham-Control infected* |  | -5.189212 | -7.525833 | -2.852590 | **< 0.001** |
|  | *Selected infected-Control sham* |  | 12.416102 | 10.079481 | 14.752724 | **< 0.001** |
|  | *Selected sham- Control sham* |  | 3.182167 | 1.018876 | 5.345458 | **0.005** |
|  | *Selected sham- Selected infected* |  | -9.233935 | -11.570557 | -6.897314 | **< 0.001** |
| **Melanization and**  **Coagulation**  **CV2** | *Control sham-Control infected* |  | -7.91494058 | -10.251562 | -5.578319 | **< 0.001** |
|  | *Selected infected-Control infected* |  | -11.26651417 | -13.764468 | -8.768561 | **< 0.001** |
|  | *Selected sham-Control infected* |  | -7.85677696 | -10.193398 | -5.520156 | **< 0.001** |
|  | *Selected infected-Control sham* |  | -3.35157358 | -5.688195 | -1.014952 | **0.006** |
|  | *Selected sham- Control sham* |  | 0.05816362 | -2.105127 | 2.221455 | 0.999 |
|  | *Selected sham- Selected infected* |  | 3.40973721 | 1.073116 | 5.746359 | **0.006** |

**Table S22: Post hoc test statistics from CVA of** **phagocytosis and encapsulation related genes.** This table shows differences between gene expression profiles of phagocytosis and encapsulation related immune genes in flies from control and selected regimes post-infection.

| **Phagocytosis and**  **encapsulation**  **CV1** | **Experimental treatments** |  | diff | lower | upper | p-adj |
| --- | --- | --- | --- | --- | --- | --- |
|  | *Control sham-Control infected* |  | -16.763209 | -19.099830 | -14.4265876 | **< 0.001** |
|  | *Selected infected-Control infected* |  | -6.062306 | -8.560259 | -3.5643524 | **< 0.001** |
|  | *Selected sham-Control infected* |  | -18.507104 | -20.843725 | -16.1704823 | **< 0.001** |
|  | *Selected infected-Control sham* |  | 10.700903 | 8.364282 | 13.0375247 | **< 0.001** |
|  | *Selected sham- Control sham* |  | -1.743895 | -3.907186 | 0.4193963 | 0.127 |
|  | *Selected sham- Selected infected* |  | -12.444798 | -14.781419 | -10.1081766 | **< 0.001** |

**Table S23: Post hoc test statistics from CVA of ROS mediated immune genes.** This table shows differences between gene expression profiles of ROS mediated immune genes in flies from control and selected regimes post-infection.

| **ROS mediated**  **CV1** | **Experimental treatments** |  | diff | lower | upper | p-adj |
| --- | --- | --- | --- | --- | --- | --- |
|  | *Control sham-Control infected* |  | 8.4308026 | 6.094181 | 10.767424 | **< 0.001** |
|  | *Selected infected-Control infected* |  | 0.8692512 | -1.628702 | 3.367205 | 0.717 |
|  | *Selected sham-Control infected* |  | 12.4111515 | 10.074530 | 14.747773 | **< 0.001** |
|  | *Selected infected-Control sham* |  | -7.5615514 | -9.898173 | -5.224930 | **< 0.001** |
|  | *Selected sham- Control sham* |  | 3.9803489 | 1.817058 | 6.143640 | **0.001** |
|  | *Selected sham- Selected infected* |  | 11.5419003 | 9.205279 | 13.878522 | **< 0.001** |

**Table S24: Post hoc test statistics from CVA of ROS mediated immune genes.** This table shows differences between gene expression profiles of ROS mediated immune genes in flies from control and selected regimes post-infection.

| **Reproduction**  **CV1** | **Experimental treatments** |  | diff | lower | upper | p-adj |
| --- | --- | --- | --- | --- | --- | --- |
|  | *Control sham-Control infected* |  | -3602154 | -3602154 | -3602154 | **< 0.001** |
|  | *Selected infected-Control infected* |  | 48981299 | 48981299 | 48981299 | **< 0.001** |
|  | *Selected sham-Control infected* |  | -1540636 | -1540636 | -1540636 | **< 0.001** |
|  | *Selected infected-Control sham* |  | 52583453 | 52583453 | 52583453 | **< 0.001** |
|  | *Selected sham- Control sham* |  | 2061518 | 2061518 | 2061518 | **< 0.001** |
|  | *Selected sham- Selected infected* |  | -50521935 | -50521935 | -50521935 | **< 0.001** |
